# Quantitative mapping of heterochrony to species-specific phenotypes

**DOI:** 10.64898/2026.06.15.732302

**Authors:** Jess J. Bourn, Simon A. Knoblich, Sophie J. Kneeshaw, Joergen Benjaminsen, Lucie Zilova, Alexander Aulehla, Lauren Saunders, Joachim Wittbrodt, Ewan Birney, Michael W. Dorrity

## Abstract

The genetic program of animal development is conserved, but its rate of execution varies across species. Heterochrony, shifts in the relative timing of developmental events, generates phenotypic variation, but its prevalence and origin are unclear. Here, we compare two vertebrates with a 3-fold difference in developmental rate, zebrafish (*Danio rerio*) and medaka (*Oryzias latipes*) and characterize heterochrony with embryo-scale single-cell genomics. We generated an atlas of >1.2M single-cell transcriptomes of medaka from blastula to hatch and developed a new approach to represent medaka development in “zebrafish time”, uncovering many cryptic, cell-level timing shifts not predicted by medaka’s slower development. We confirm a divergent heterochrony in the medaka notochord using in vivo imaging, revealing that coupled acceleration and delay of sister cell types shapes the species-specific morphology of this tissue. Our results point to cell type-specific timing deviations as a reservoir of phenotypic variation and we propose that species-specific developmental rate can emerge from these cell-level differences.

## Introduction

Embryonic development unfolds via sequential activation of a conserved genetic program that triggers emergence of distinct cell types (*1*, *2*). The rate at which this program unfolds defines organism-level developmental rate, or tempo, which varies across species (*3*). The presence of widespread tempo variation despite conservation of developmental genes has led to a search for global regulators eliciting proportional shifts in the timing of cell differentiation across species (*4–7*). However, it remains unclear whether tempo is directed at the organism-level or if it emerges from underlying cells or tissues. Without identifying the source of developmental tempo, we cannot define timing control mechanisms or predict the phenotypic consequences of developmental timing variation (*8*, *9*). For example, increasing temperature accelerates tempo but challenges coordinated differentiation at the cell-level within-species (*10*), while even stronger cell-level shifts occur naturally between species (*11*, *12*). The latter phenomenon, termed “heterochrony,” is defined as a relative change in the rate, timing, or order of cellular or developmental events (*12*, *13*). Because heterochrony occurs as a disproportionate shift at the cell- or tissue-level relative to organism-level developmental progression, it serves as a source of phenotypic variation among species that share conserved developmental genes and cell identities (*14–16*).

Classically, heterochrony has been defined morphologically, limiting its detection to cases where both organism-level progression and an underlying cell or tissue-level shift can be clearly defined between species (*13*), but single-cell genomics has enabled alignment of developmental events such as differentiation and maturation of cells (*17*). Accordingly, cross-species comparisons of both embryos and *in vitro* tissue models have revealed abundant timing variation (*4*, *6*, *7*, *17–22*). Advances in sample multiplexing have further increased sensitivity by estimating developmental age for hundreds of embryos and constituent cells (*10*, *23–26*) in a single experiment, capturing developmental timing variation across individuals. However, without systematic, generalizable tools for disentangling heterochrony from organism-level scaling of developmental processes, current approaches are unable to resolve whether timing control mechanisms operate at the organism or cellular level and how they influence species-specific phenotypes.

In this study, we sought to systematically characterize heterochrony in vertebrate development, resolving how timing differences map to tissue- and organism-level phenotypes of teleost fish, a diverse group with conserved embryonic patterning (*27*). The model teleosts medaka (*Oryzias latipes*) and zebrafish (*Danio rerio*) share more than half of their genes as 1-to-1 orthologs (*28*, *29*), yet their developmental tempo differs by two- to three-fold. This divergence is remarkable given that both species have similar thermal optima and adult body size (*30*), traits which are commonly linked to developmental tempo (*8*, *31*). We construct a time-resolved developmental atlas of medaka from blastula to hatch, comprising more than 1.2 million cells from over 500 samples to enable comprehensive cross-species comparisons of developmental timing at both embryo and cell levels. We integrate these data with an embryo-resolved atlas of zebrafish to place both organisms along a shared, transcriptionally-defined developmental trajectory, collapsing clock-time differences in their developmental tempo. Within this framework, we introduce a generalizable measure, “cell-embryo deviation,” to quantitatively capture heterochronic shifts across cell types between species. In visualizing the cellular consequences of a divergent heterochrony within the notochord, we show how timing shifts within a tissue influence a medaka-specific phenotype. Using our heterochrony metric for unbiased detection of genes associated with timing, we find that developmental timing differences are not governed by a small set of global regulators, but instead via cell type-specific combinations of molecular processes. Together, our results indicate that developmental timing is primarily controlled at the cell type-level, rather than by a single, embryo-wide clock mechanism.

## Results

### A comparative resource for identifying cell type-specific timing differences

To capture temporal variation in medaka development, we performed optimized, combinatorial indexing-based single-nucleus RNA sequencing (sci-RNA-seq3, see Supplementary Text) (*24*) on 533 medaka Cab embryos sampled over 19 developmental timepoints, spanning early blastula to late hatching stages (Fig. 1A). We used hash barcoding by sci-Plex to sample 24 embryos per timepoint, capturing variation in developmental events across embryos. Quality control and doublet filtering resulted in a dataset consisting of 1,156,489 cells with an average of 2194 cells captured per embryo (Fig. S1A). We performed unsupervised clustering to group cells and cluster-specific marker genes were then used to assign annotations at three levels (*32*, *33*). We first grouped cells into 14 “major groups”, most of which represented tissues (e.g. muscle, retina, liver) over all timepoints; a single “major group” with cells from the earliest timepoints (6.5 hpf to 26 hpf) represented the blastula to late neurula stages. Major groups were processed and annotated separately at finer resolution. In total, we hierarchically classified all cells in the dataset into 39 tissue types, 198 broad cell types and 302 sub cell types (Figure 1B, Table S1), capturing the developmental trajectories of all major cell lineages during medaka embryogenesis. To confirm that defined cell types were stable and validate use of these data as a reference, we sampled 24 embryos from diverse medaka strains (Kaga, HNI, MIKK72 and MIKK79) at timepoints spanning the range of our atlas (*34*, *35*). Co-embedding of strain data with the Cab reference revealed coherent mapping of cell type and timepoint labels across datasets, despite batch and genetic differences (Fig. S1B).

**Figure 1.**
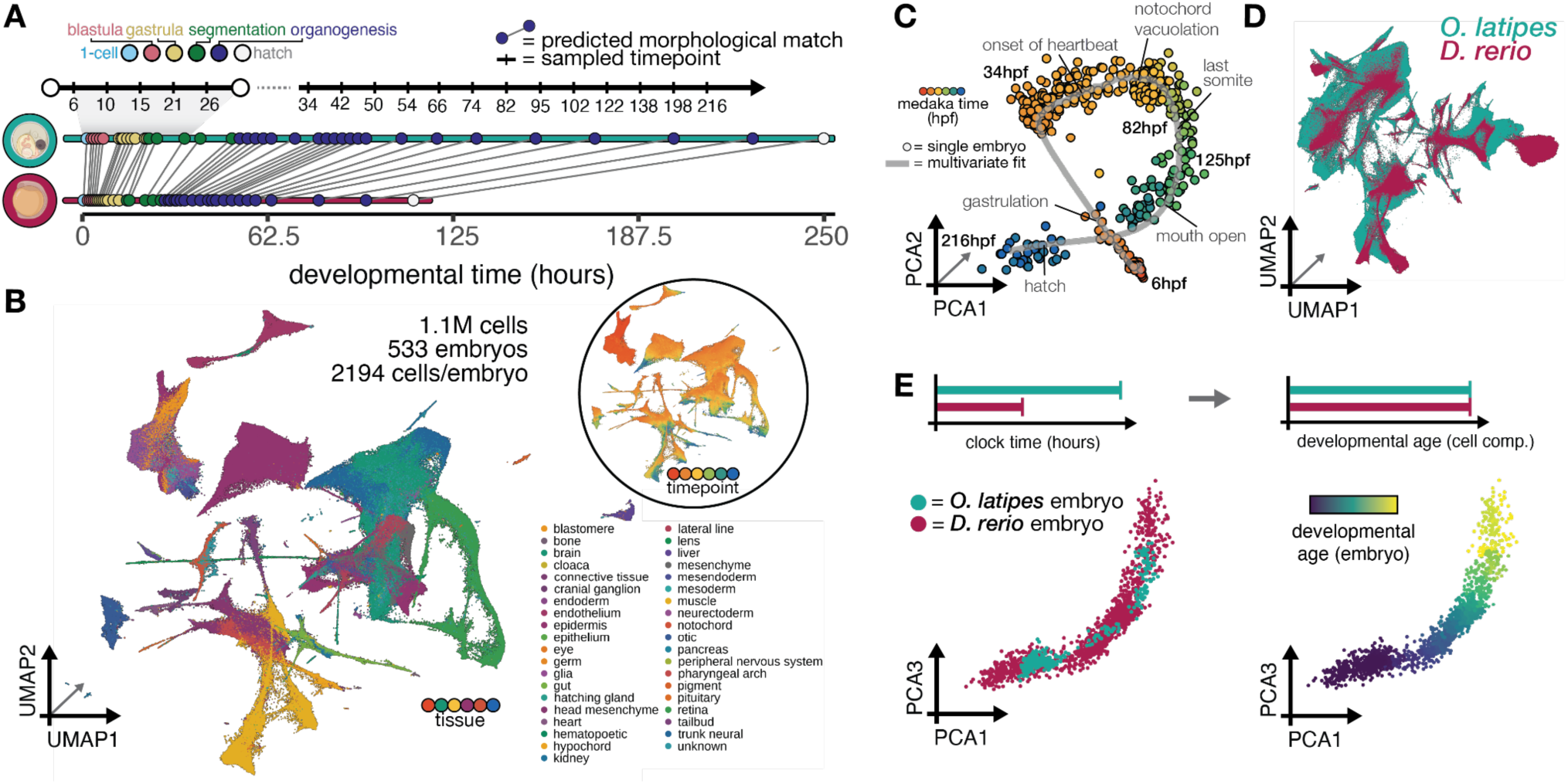
A time-resolved atlas of medaka development from blastula to hatch. (A) Schematic overview of sampling design for medaka sci-RNA-seq3 atlas, illustrating the developmental timepoints sampled from 6.5 to 216 hours post fertilization (hpf), and their corresponding stages represented in zebrafish based on morphological staging (*28*). (B) Uniform manifold approximation and projection (UMAP) of the medaka atlas embedded in three dimensions, colored by tissue annotation. Inset colored by per-cell developmental age. (C) The embryo-level trajectory of medaka development is a simple curve through the first three principal components of cell composition space; each point represents a single sampled embryo and is colored by clock timepoint. Cell compositions predicted from a multivariate fit as a function of timepoint are shown in grey. (D) Medaka (teal) and zebrafish (maroon) time-series data represented in a shared UMAP coordinate space in three dimensions, constructed by MNN-correction for species on a shared feature set of 1:1 orthologous genes. (E) Embryo trajectory constructed using cell composition per embryo, as in (C), but using, for both species, shared cell type labels from zebrafish, colored by species (left) and per-embryo pseudostage values (right).

We first sought to define embryo-level developmental progression in medaka. We reasoned that, independent of the clock-time duration of embryogenesis, progression is fundamentally defined by the coordinated gain and loss of cell identities over time. We therefore quantified how the abundance of each cell type, and the overall cellular composition of the embryo, changed over the course of embryogenesis. For each individually barcoded embryo, we aggregated cells of each broad cell type to generate a cell type *x* embryo matrix describing the cell type composition for every medaka sample. We analyzed cell type complexity over the course of embryogenesis and found that the number of unique cell types sampled per embryo saturates despite an increasing number of total cells per embryo over time (Fig. S1, C to E). We then used principal component analysis (PCA) to examine per-embryo cell composition in low dimensional space (Fig. 1C). The samples fell on a simple, one-dimensional trajectory capturing the ordered progression of development; a continuous, quantitative analog to the embryological definition of developmental progression as movement through a discrete set of “stages” indicated by whole-embryo morphology. We defined an underlying cell composition trajectory using a multinomial regression of all cell counts over time, and assigned individual medaka embryos a “pseudostage” (*10*) using their position on this trajectory. Thus, embryo-level developmental time can be inferred from cell composition data, enabling clock-time independent embryo staging and capturing stage variation even within sample timepoints.

This finding prompted us to ask whether the same approach could be applied to define a clock-time independent method of cross-species stage matching to zebrafish. Classical stage matching relies on morphological criteria that, although diagnostic of developmental progression, can be challenging to quantify and to compare across species (*22*, *28*, *36*). Using a previously-generated sci-Plex zebrafish atlas (*25*), we co-embedded cells of overlapping developmental stages from both species using only conserved 1-to-1 orthologs (n = 15,011) as gene features (Fig. 1D) (*37*). Using cell clusters defined in the shared coordinate space, we computed a joint cell type composition matrix containing both zebrafish and medaka embryos. We validated the shared coordinate space by comparing our manual medaka annotations to those predicted by transferring cell type labels from zebrafish, finding 79% of these matched across label sets (Fig. S1F). We performed PCA analysis of the joint cell composition matrix and found that zebrafish and medaka embryos fall along a shared trajectory in low dimensional space (Fig. 1E, left panel), indicating an overall conserved developmental progression at the embryo level, despite differences in their clock time duration. We assigned “pseudostage” values in the joint cell composition space to capture clocktime-independent stages for all embryos (Fig. 1E, right), serving as the first continuous stage alignment between zebrafish and medaka (Fig. S1G).

### Species-specific elaboration of a conserved cellular identity

While our cross-species pseudostaging approach captures broad, conserved developmental processes between medaka and zebrafish at the embryo-level, it may also obscure species-specific differences at the cell-level. To identify medaka-specific cell types, we next analyzed the medaka data alone, as the cross-species co-embedding depends on a shared feature set of only 15,011 1-to-1 orthologous genes (∼⅔ total medaka genes; Table S2). We focused on a medaka-specific cell population associated with rhythmic contractile waves during early medaka embryogenesis, reported to originate in a layer of so-called “stellate cells” (*38*). Zebrafish lack such contractions, despite showing similar morphology and patterning during early development. To identify putative stellate cells, we analyzed medaka sci-RNA-seq data from blastula up to early segmentation (Fig. 2A), hypothesizing that, despite occupying a similar trajectory with zebrafish at the embryo-level, medaka may show functional divergence at the cell-level. Using live imaging, we confirmed the presence of rhythmic contractions in medaka Cab embryos, which emanate ventrally and span the ventral two-thirds of the embryo (Fig. 2B, movie S1). These contractions arise in medaka (Iwamatsu stage 14, 15 hpf) (*39*) during the zebrafish equivalent of 50% epiboly to shield stages (Kimmel stage 4, 5 hpf) (*28*, *40*). We next sought to identify the transcriptional identity of cells using prior descriptions of calcium-dependence in these cells as a guide (*41*, *42*). We identified a putative population of stellate cells at developmental timepoints coinciding with the onset of the rhythmic contractions (Fig. 2C, S2A). These cells expressed conserved markers of the non-neural ectoderm and its epidermal derivatives (*tp63*, *gata2a*, *dlx3b*, *tfap2c*), and were transcriptionally distinct from the cells of the enveloping layer (EVL; *krt4+, ace+*) (Fig. 2C).

**Figure 2.**
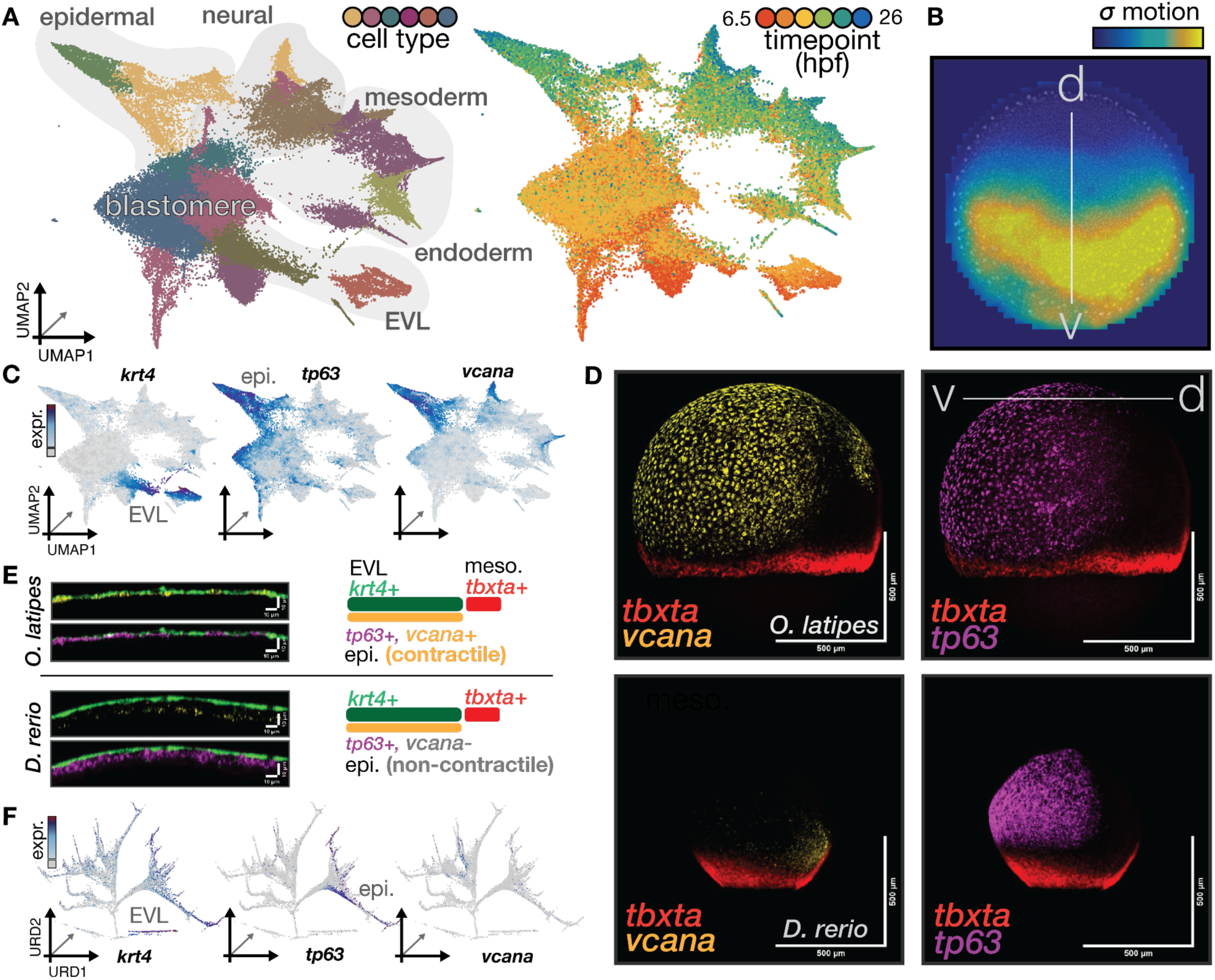
A medaka-specific cell type shows early activation of contractility genes in a conserved cell identity. (A) UMAP of early medaka cells (6.5 to 26 hpf, spanning blastula to late epiboly stages) embedded in three dimensions, colored by broad cell type annotation (left) and timepoint (right). (B) Quantification of rhythmic contractile activity in the medaka embryo during gastrulation stages using particle image velocimetry (PIV). Colors indicate the standard deviation of velocity magnitude (*σ* motion), which is confined to the ventral two thirds of the blastoderm (dorsal-ventral axis indicated). (C) UMAP plots of early medaka cells colored by log10 expression of *krt4* (EVL), *tp63* (epidermal), and *vcana* (stellate cells). (D) Whole-mount hybridization chain reaction (HCR) staining of a 25 hpf (Iwamatsu stage 17) medaka embryo (top row), showing the spatial distributions of *tbxta* (red) and *tp63* (purple) expression, and confirming *vcana* (yellow) expression in the contractile ventral domain. HCR for identically oriented 9 hpf (Kimmel 90% epiboly) zebrafish embryos show absence of *vcana* in the ventral domain (bottom row). (E) Orthogonal view of HCR stainings in D shows spatial separation of the epidermal (*tp63*, purple) and EVL (*krt4*, green) layers in both species, with expression of *vcana* in the epidermal layer of medaka (top) and no expression in either layer in zebrafish (bottom). (F) Reduced dimension (URD) plots of early zebrafish cells (*32*) colored by log10 expression markers shown in (C); no *vcana* expression is observed in epidermal cells in zebrafish.

To confirm that this population corresponded to a spatially identifiable cell type *in vivo*, we performed hybridization chain reaction (HCR) on morphologically stage-matched medaka and zebrafish embryos; *tp63* (epidermis) and *krt4* (EVL) showed near identical expression in ectoderm in both species (Fig. 2, D and E, Fig. S2B), suggesting a conserved cell identity and embryonic organization. However, this population shows a surprising, medaka-specific molecular signature; *vcana,* a proteoglycan expressed later in the developing heart, was robustly expressed within the medaka *tp63*+ stellate cell population, while *vcana* expression was not detected in the comparable *tp63*+ epidermal population of zebrafish embryos (Fig. 2, D and E). The large number and organization of *vcana*-expressing cells in medaka led us to hypothesize that stellate cells do not constitute a novel cell identity, but instead represent a contractile variant of a conserved epidermal cell population (Fig. S2, C and D). Indeed, we validated early epidermal expression of genes enriched for extracellular matrix (ECM) components and calcium signaling pathways (*vcana, fbn2b, hapln3, hmcn2, actin-pls3* and *cacna1c)* in medaka (Fig. S2E). Many of these genes are associated with calcium-mediated contractility and are strongly conserved, but are not expressed in the corresponding epidermal population (*tp63+, dlx3b+, tfap2c+)* in zebrafish (Fig. 2F, Fig. S2F) (*32*). Both the single-nucleus and *in vivo* results indicate that the medaka-specific function of stellate cells is built on top of a core epidermal cell identity that is recognizable between species. Genes associated with contractility in medaka are 1-to-1 orthologs that are also co-expressed in zebrafish, albeit in cell populations arising later in development (*32*, *43*). Thus, we next used our comparative framework to systematically analyze variation in the timing of developmental events.

### Systematic quantification of heterochronic shifts between species

While medaka embryos could be pseudostaged (*10*) and aligned with zebrafish (Fig. 1E), we wondered whether cell-level timing relationships might differ between species, with specific cell types occurring earlier or later relative to the overall embryo stage alignment. Thus, we sought to represent developmental progression at both the cell type-level and the embryo-level. We estimated cell-level developmental age by averaging, for each cell, the timepoints of its nearest transcriptional neighbors in the medaka time-series atlas, providing a quantitative metric with units in medaka hours (Fig. S3A). We estimated embryo-level developmental age by taking a simple average of the estimated age for all cells derived from the same embryo. Embryo-level age estimates were strongly correlated with “pseudostage” values from the cell composition-based embryo trajectory and captured substantial within-timepoint variation in embryo stage (Fig. S3B). We next put these estimates into a cross-species context, asking how cell-level age deviates from the embryo-level age in medaka vs zebrafish. For simplicity, we examined medaka progression relative to the faster pace of zebrafish development. We used the cross-species co-embedding to transfer zebrafish timepoint labels to every medaka cell in the shared coordinate space, matching species-specific time based on transcriptional similarity (Fig. S3C). We then used the above procedure to compute embryo- and cell-level age, resulting in embryo- and cell-level representations of medaka development in within-species “medaka time” and cross-species “zebrafish time” (Fig. 3A, Fig. S3C).

**Figure 3.**
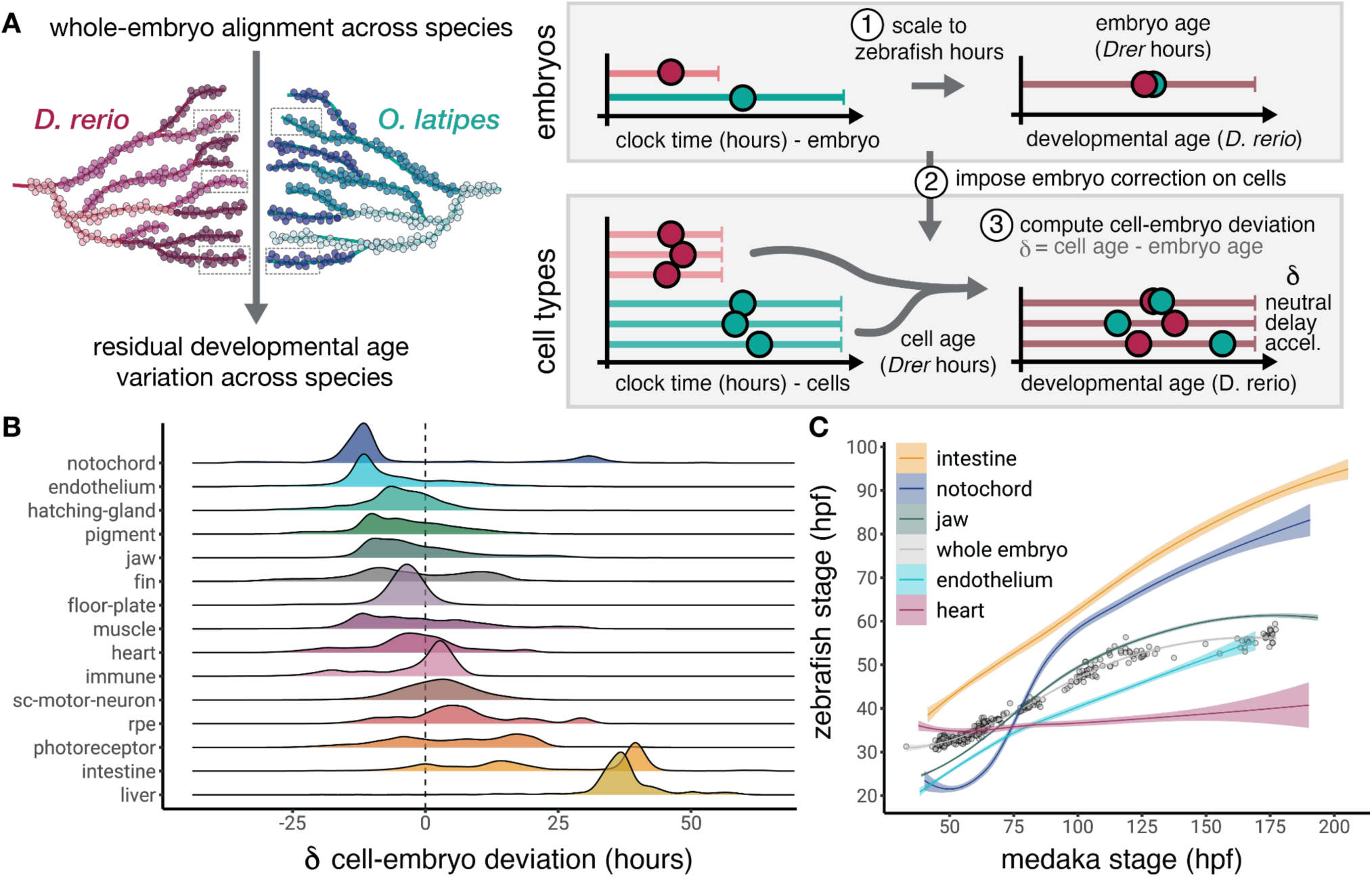
Systematic quantification of cell type heterochrony. (A) Schematic for quantifying cell-typespecific heterochrony. Whole-embryo developmental progression is first aligned between zebrafish (*D. rerio*) and medaka (*O. latipes*) using embryo-level transcriptional similarity (left). In parallel, cell types are aligned across species based on cell-level transcriptional similarities (right). Both values are computed in zebrafish hours. For every cell, the cell-embryo deviation (CED) is calculated as the difference between its cell-type-specific developmental age and the corresponding embryo-level developmental age for the individual animal the cell was derived from. Positive values indicate that a cell type shows a relative acceleration in development compared to embryo-wide progression of zebrafish, whereas negative values indicate relative delay. (B) Distributions of CED values across multiple medaka cell type trajectories, ordered by increasing average CED. (C) Cross-species developmental trajectories for whole embryo and individual cell types, showing predicted zebrafish age (hpf) as a function of medaka age (hpf). Lines represent smoothed fits for whole-embryo alignment (grey) and individual cell types (colors), built on data where each point represents the average age of that cell type in each embryo (n = 367). Shaded regions indicate the 95% confidence interval.

When comparing cell-level and embryo-level ages in “zebrafish time,” we found that, rather than showing a consistent match, medaka cells were often assigned ages that were remarkably different from the assigned embryo-level age, suggesting cell-level differences that are not captured in whole-embryo comparisons. To analyze cross-species age discrepancies across all medaka cell types, we quantified the difference in cell-level and embryo-level age in zebrafish hours. We termed this quantity “cell-embryo deviation” (δ or CED), which denotes the amount by which a specific medaka cell or cell type is accelerated or delayed in its developmental progression when compared to zebrafish (δ_medaka cell_ = cell age*^Drer^* (hpf) *-* embryo age*^Drer^* (hpf)) (Fig. 3A). When computed within-species, the CED metric captures natural timing variation in cell- vs embryo-level progression across individual animals. We first validated the CED approach to detect cell-level timing deviations in the within-species context by analyzing sci-Plex data for zebrafish mutants of the transcription factor *tbx16*, whose embryos look morphologically ‘normal’ in the anterior and severely delayed in the posterior due to arrest of mesoderm progenitors (*25*, *44*, *45*). In this scenario, we expect cell-level delay (CED < 0) in the mesoderm, while other cell types should progress with the embryo-level age (CED near 0). Using the CED approach, we find that mesodermally-derived cell types are developmentally “younger” than expected of the embryo as a whole (Fig. S3D) and that the embryo overall shows a developmental age matching the clock-time of the sample (average age of mutant embryos sampled at 36 hpf = 36.6 hpf), owing to the majority of cells progressing as normal. Thus, CED captures the delay of the affected mesodermal cells and the normal progression of other cells in the embryo, disentangling cell-level timing variation from embryo-level progression (*10*, *25*, *46*).

We next applied CED to understand the discrepancies in cellular and embryo age in the cross-species context. We selected 15 functionally comparable cell types in both medaka (Fig. S3E) and zebrafish, extracted the full developmental trajectory (all timepoints) of each from the two sci-Plex datasets, and computed CED for all cells. Some trajectories, such as the heart or motor neurons, showed CED around 0 and scale their progression with the embryo as a whole. However, we observed clear biases in CED distribution across most cell types, suggesting that deviations from embryo-level tempo, rather than being an exception, are a common feature of medaka development relative to zebrafish (Fig. 3B). Several trajectories, such as notochord and endothelium, showed predominantly negative CED values, indicating that these medaka cells are more transcriptionally similar to earlier zebrafish stages and are thus relatively delayed in medaka. In contrast, other trajectories, such as the liver, displayed strongly positive CED, suggesting similarity to later zebrafish developmental timepoints and accelerated progression in medaka, a heterochrony that has been previously suggested based on morphological comparison (*36*). Many cell types showed broad or multimodal distributions; pigment, fin and photoreceptor trajectories exhibited wide distributions spanning both negative and positive CED, suggesting complex temporal reorganization in these tissues relative to zebrafish. To ensure cross-species CED variation was not due to technical bias, we computed a null distribution derived from natural timing variation of these cell types within-species (Fig. S3F). Within-species CED distributions were centered close to zero for both zebrafish and medaka and were not globally offset. Cross-species CED distributions in medaka frequently exceed this within-species expectation and likely reflect genuine heterochrony. Despite their morphological similarity, embryo-level comparisons alone are not sufficient to capture the extent of developmental timing differences between medaka and zebrafish; many cells deviate from the expectation of a global slowing of development in medaka.

Finally, we asked whether cell type-specific timing variation uncovered with our approach represented linear acceleration or delay or if cells instead become uncoupled from the global tempo of the organism at different stages of development. When represented in zebrafish age, medaka cell types displayed individual and distinct dynamics over the span of embryogenesis (Fig. 3C, Fig. S3G), with many showing residual variation beyond the expectation from the whole-embryo relationship between species. For example, the medaka liver shows a relatively earlier emergence time and a positive residual over the course of development, revealing advanced progression that is maintained throughout medaka stages. In contrast, the notochord displays a change in the sign of this residual over time, shifting from negative to positive as development proceeds. All other cell types showed distinct patterns throughout medaka embryogenesis and were remarkably specific in their dynamic; some revealed initiation or maturation shifts (fin), overall accelerated or delayed rate differences (liver, pigment), or both (notochord). Thus, our approach can be used to identify shifts in emergence, duration or rate of a developmental process, with time resolution of even a few hours. However, the observation of distinct dynamics across cells in medaka suggests that a global, embryo-wide tempo mechanism does proportionally slow development at the cell level.

### Species-specific phenotype driven by divergent heterochrony within the notochord

In the apparent absence of global coordination across the embryo, we next asked whether timing differences were coordinated at the tissue level. Guided by our heterochrony analysis, we focused on the notochord, a tissue composed of two morphologically and functionally distinct cell types, inner vacuolated cells and an outer layer of sheath cells, derived from a common progenitor, and whose markers are conserved between zebrafish and medaka (Fig. 4A, Fig. S4A). The medaka notochord showed a divergent trend in CED, with some cells delayed and some accelerated, shifts we verified by computing false discovery rates for across- vs within-species CED of notochord development (Fig. S4B). The population exhibiting relative delay in development contained notochord progenitors and vacuolated cells from earlier stage embryos, while the accelerated population contained sheath cells from later stage embryos (Fig. 4, B and C, Fig. S4C). To identify genes driving species-specific temporal shifts in notochord development, we examined cross-species temporal correlation of 1-to-1 orthologs over the shared developmental trajectory (Fig. S4D, Table S3), and visualized their aggregated expression in each cell population. Genes with temporally conserved expression across species (*emilin3a*, *tbx3a*, *slit3* and *ephb3*) were strongly enriched in the progenitor cell population (Fig. S4E), while genes with the lowest correlations were expressed in vacuolated and sheath cells (Fig. 4D). This trend was consistent across all notochord genes (Fig. S4F), suggesting that divergent shifts in developmental timing are possible even within a tissue whose cells share a progenitor pool.

**Figure 4.**
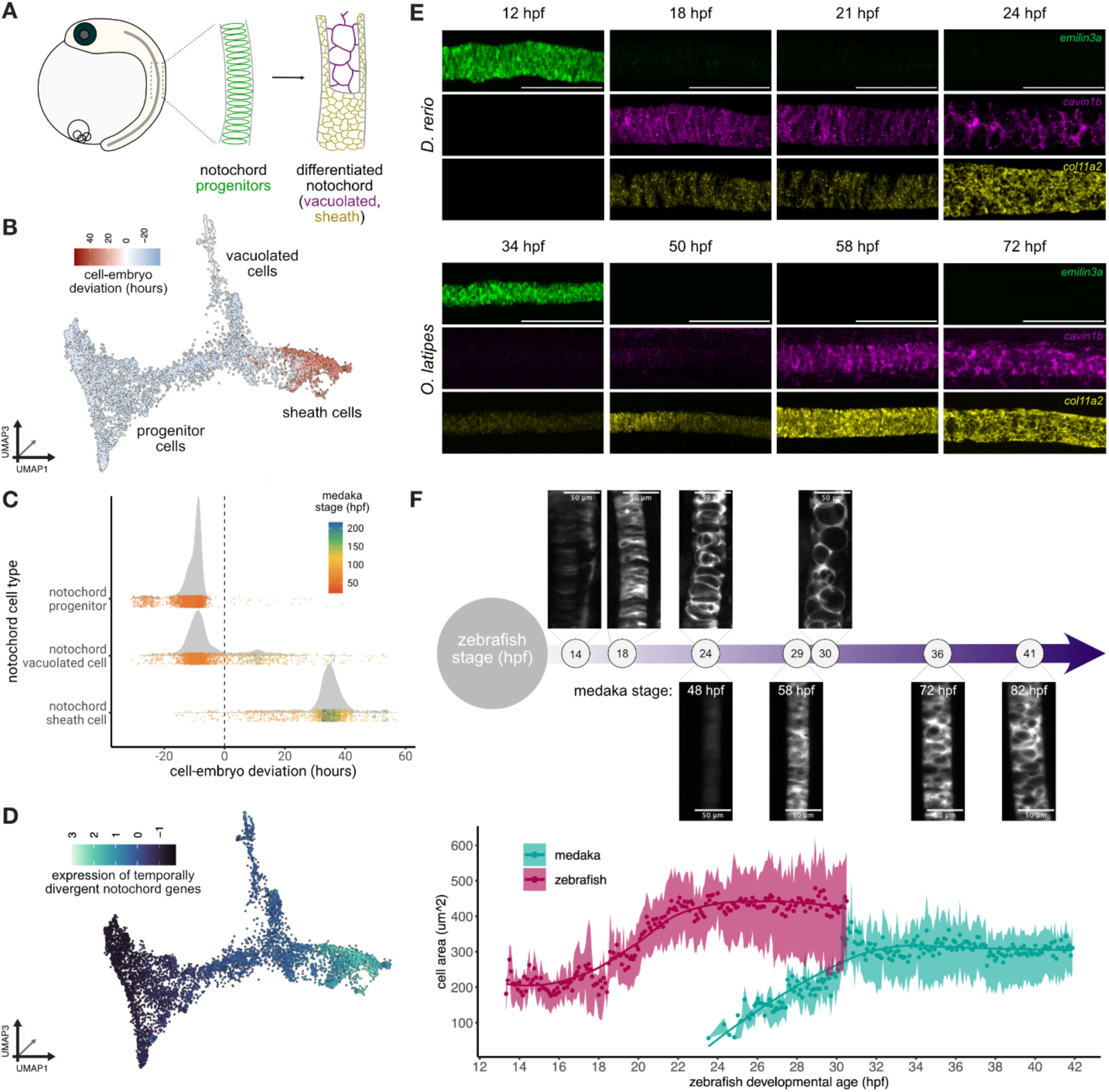
Notochord differentiation dynamics are reorganized in medaka to accommodate slower development. (A) Schematic illustrating conserved notochord differentiation dynamics in medaka. The disc-shaped progenitors give rise to two populations of mature cells, the large inner vacuolated cells and the surrounding layer of smaller sheath cells. (B) UMAP of medaka notochord cells colored by CED, indicating relatively delayed (progenitor, vacuolated) and accelerated (sheath) cell populations. (C) Distributions of CED values for each cell population in the medaka notochord, with distributions of cellular age shown beneath. (D) The aggregated expression of the 1:1 orthologous genes with the most divergent temporal dynamics across species; sheath cells display the highest expression levels of temporally divergent genes. (E) Time-series of hybridisation chain reaction (HCR) stainings illustrating the progression of notochord cell differentiation at morphologically stage-matched timepoints in zebrafish (top) and medaka (bottom). Within species, each panel shows progenitor (green, *emilin3a*), vacuolated (magenta, *cavin1b*) and sheath (yellow, *col11a2*) cells, revealing species-specific timing differences in the onset and progression of notochord differentiation. Scale bars = 100 μm. (F) Live imaging of notochord vacuolation dynamics in Tg(*jag1a::mNeonGreen*) zebrafish and Tg(*desmog::eGFP*) medaka embryos, aligned by morphological stage. Top, representative images of notochord vacuolation dynamics in zebrafish (top row) and medaka (bottom row), with clocktime developmental age indicated. Bottom, quantification of average vacuolated cell area over time plotted against stage-matched zebrafish developmental age. Points represent average cell size at each timepoint, lines indicate smoothed fits over all frames in the movie, and shaded regions show interquartile range of cell size per timepoint for n = 3 samples of each species. Medaka vacuolated cells begin expansion later than zebrafish cells despite temporal alignment, indicating delayed morphological progression. Scale bars = 50 μm.

To test whether these transcriptionally-inferred timing shifts have cellular, morphological, or functional consequences in the native embryo context, we characterized this heterochrony *in vivo*, combining *in situ* and live imaging in stage-matched embryos of both species. We first targeted conserved 1-to-1 orthologs specific for the progenitor cells (*emilin3a+*), vacuolated cells (*cavin1b+*) and sheath cells (*col11a2+*), and visualized their expression with HCR (Fig. 4E, Fig. S4, G and H). At the earliest timepoint (6-somite stage), *emilin3a* displayed a similar expression pattern in both species, supporting temporal conservation of expression in progenitor cells (Fig. 4E, leftmost panels). The next stage-matched timepoint (18-somite stage) showed a clear, species-specific reversal in the order of cell maturation, with zebrafish notochords dominated by *cavin1b* expression and moderate *col11a2* signal, while medaka notochords showed *col11a2* expression and little detectable *cavin1b* (Fig. 4E, middle-left panels). At later stages, expression patterns were more similar, with both species containing clear populations of sheath and vacuolated cells, although medaka showed continued enhanced *col11a2* expression relative to zebrafish (Fig. 4E, middle-right panels). Thus, *in situ* imaging recapitulates the divergent heterochrony predicted by CED. Consistent with this pattern, we used per embryo cell counts to show that differential maturation dynamics of sheath and vacuolated cells results in endpoint differences in their proportions that reflect species-specific notochord morphology (Fig. S4I) (*47*, *48*).

To determine whether these transcriptional shifts correspond to altered cellular function, we next examined notochord vacuolation dynamics using live imaging. In both species, vacuolated cell maturation rate has the potential to affect notochord morphogenesis; as these cells progressively expand their intracellular vacuoles, they form larger, more rigid cells that contribute to physical properties of the notochord and to axis elongation (*48*, *49*). We captured the onset and progression of vacuolation in multiple embryos of each species, starting from 6-somite stage in each species (zebrafish 12 hpf, medaka 34 hpf; movie S2). We matched medaka timepoints to their zebrafish developmental equivalent using prior morphological comparisons (*36*) (Fig. S4J), enabling quantitative comparisons on a shared timescale (Fig. 4F, top). Measuring average vacuolated-cell area over time in each series, we found that medaka vacuolated cells delay their expansion relative to zebrafish (Fig. 4F, bottom), which results in an altered notochord phenotype. Fully vacuolated cells (after the 30-somite stage) in medaka have a mean diameter of 19.8 µm, 16% smaller than the diameter of mature vacuolated cells in zebrafish (23.5 µm). However, the rate of expansion (*V_max_*) is similar in both species (39.3 ± 1 µm^2^/hour for zebrafish, 36.8 ± 1.3 µm^2^/hour for medaka), suggesting that the function enacted by vacuolation genes does not directly scale with embryo-level tempo, and that observed phenotypic differences result instead from the shifted emergence time of these cells in medaka. We conclude that notochord timing differences captured by CED correspond to genuine delays in morphogenetic progression, rather than differences in gene expression alone. Together, these *in vivo* data support the conclusion that although medaka develop slower than zebrafish, the developmental rate of individual cell types do not scale uniformly with whole-embryo progression, or even within a tissue. Rather than resulting from organism- or tissue-level enforcement, developmental timing variation may instead be constrained by cell-type-specific functions such as notochord vacuolation, necessitating shifts in cellular emergence or differentiation time to enact morphogenetic changes relative to the surrounding embryo.

### Molecular processes driving heterochronic shifts are cell type-specific

To resolve the relationship between cell type-specific functions and developmental timing differences across the embryo, we next sought to systematically identify molecular processes associated with accelerated or delayed development. Using a standard regression framework, we used CED to identify genes whose expression is associated with relative acceleration or delay across cells. We performed differential gene expression analysis across 15 cell type trajectories, posing two main questions: (1) what genes are associated with global acceleration or delay across all cells; and (2) what genes are associated with local heterochronic shifts within cell types?

A total of 3576 genes were identified as being significantly (q-value < 0.05) associated with CED across cell types (Table S4), with similar numbers of genes associated with acceleration (46%) and delay (54%). Genes associated with accelerated development (positive heterochrony) were functionally enriched for terms relating to cell-cell interactions and cell migration (adhesion, actin binding and transmembrane transport), suggesting a role for cellular interactions in accelerating developmental processes in medaka (Fig. 5A, left). Genes associated with delayed development (negative heterochrony) were enriched for terms associated with RNA processing, translation, and nuclear organization (Fig. 5A, right), consistent with previous comparative studies showing that gene expression machinery and protein stability is linked to species-specific developmental tempo (*5*, *19*).

**Figure 5.**
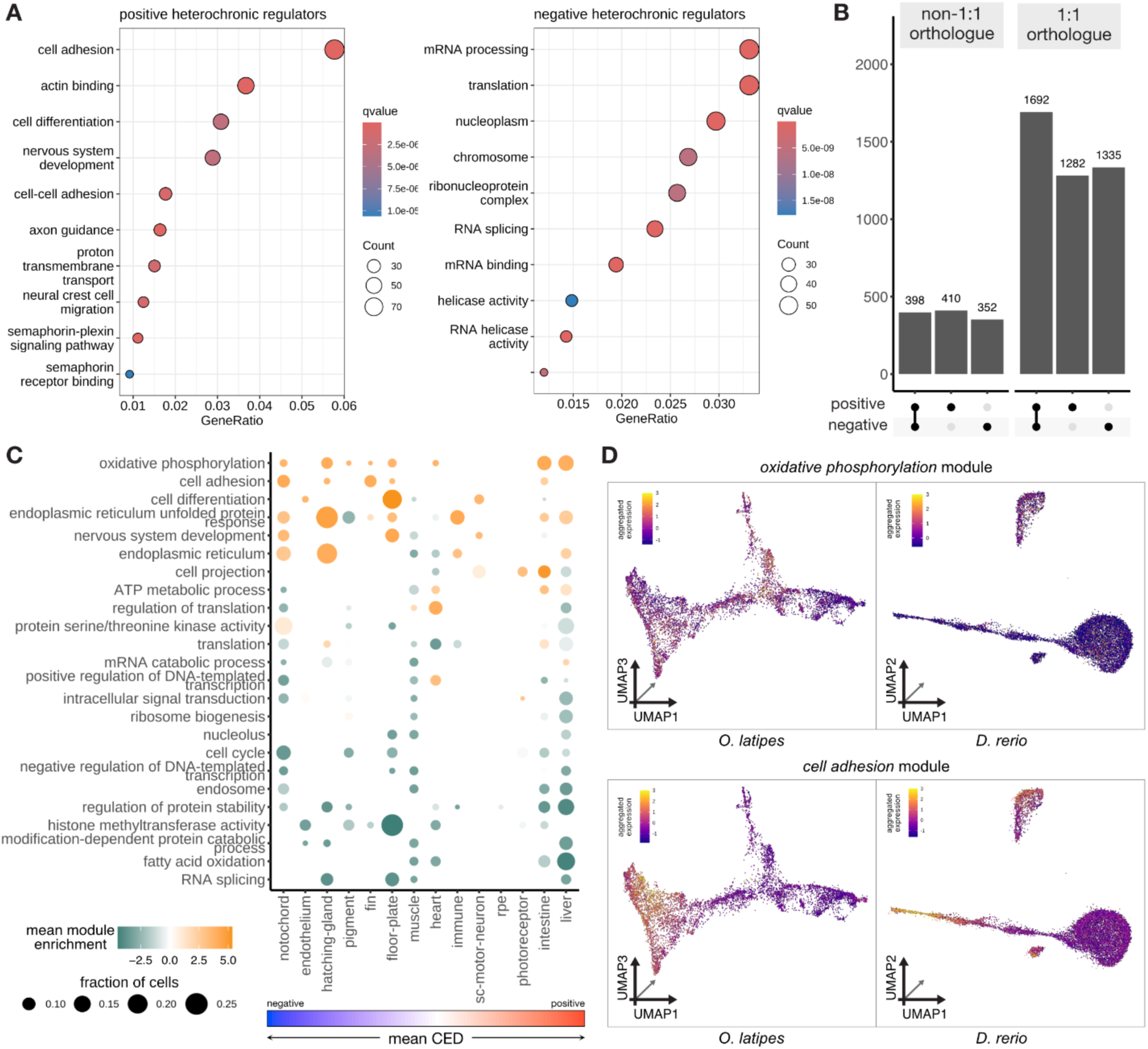
Diverse mechanisms underlie cell type-specific timing across species. (A) Gene Ontology (GO) enrichment analysis of genes significantly associated with accelerated (left; positive CED) or delayed (right; negative CED) developmental timing across cell types. Dots represent enriched GO terms, positioned by gene ratio, sized by the number of genes contributing to the term, and colored by adjusted q-value. (B) Counts of CED-associated genes separated by direction of effect (positive, negative, or bidirectional) and orthology group, showing that both accelerating and delaying regulators are strongly enriched for conserved 1:1 orthologs relative to non-1:1 genes. (C) Summary of each module’s activity across trajectories of cells above the two standard deviation threshold. Dot size indicates the fraction of cells within each trajectory showing significant module enrichment, while color shows mean module enrichment score. Trajectories (columns) are ordered by increasing mean CED and modules (rows) are ordered by average module score weighted by fraction of cells. (D) UMAP plots of the notochord trajectories in medaka and zebrafish, embedded in three dimensions. Colors represent the aggregated gene expression of the 1:1 orthologs in the oxidative phosphorylation (top) and cell adhesion (bottom) modules, illustrating both species- and cell type-specific usage of conserved molecular pathways.

We next examined genes associated with CED at the level of cell type. We identified 5469 unique genes significantly associated with CED (Table S5), many of which are reused across cell types, resulting in a total of 15,810 associations. Rather than being driven by species-specific genes, the large majority of the CED-associated genes (79%) are conserved 1-to-1 orthologs (Fig. 5B), which only constitute 62% of the medaka protein-coding genome (15,011 out of 24,316 genes). Most of these genes were associated with both positive and negative CED values in different cell types (Fig. S5A), suggesting that developmental rate control may not be governed by genes with strictly unidirectional effects.

To move from gene-level associations to molecular processes underlying heterochrony, we next examined the activity of defined gene modules and their relationship to cellular CED scores. We assembled gene modules using data-driven (Fig. 5A, Fig. S5A), and literature-driven hypotheses of developmental timing regulation (e.g., protein stability, mRNA processing, metabolism) and computed module scores for all cells across the 15 trajectories (*50*). This method correctly recapitulated cell types displaying enrichment of specific modules (Fig. S5B) and related gene module activity to CED across trajectories in accelerated or delayed developmental contexts.

Few modules showed a consistent, linear relationship with CED across cell types, but activity of ATP metabolic processes, synapse, glycolysis and ATP hydrolysis activity often increased or decreased unidirectionally with CED (Fig. 5C, Fig. S5C). Several metabolism-and epigenetic-associated modules appeared repeatedly across cell types, suggesting that these processes commonly contribute to developmental rate control, even if their directionality varies. No module was uniformly up- or down-regulated across all accelerated or all delayed cell types, suggesting that no single pathway globally directs developmental tempo differences between zebrafish and medaka. Instead, many modules were active in both contexts (bidirectional), implying that the functions of these modules may be employed differently depending on cellular identity. Certain modules, such as oxidative phosphorylation, regulation of protein stability, and histone methyltransferase activity, were enriched unidirectionally across cell types with markedly different CED values (Fig. 5C). For example, the oxidative phosphorylation module was upregulated in the notochord, pigment, floor plate and liver, while the module for regulation of protein stability was consistently downregulated in the notochord, hatching gland, intestine and liver, despite these cell types spanning strongly negative to strongly positive CED values. We examined two gene modules, oxidative phosphorylation and cell adhesion, in the context of the divergent heterochrony we identified in the medaka notochord, finding elevated expression of both modules in the delayed notochord progenitors, consistent with their negative CED (Fig. 5, C and D). The oxidative phosphorylation module was not enriched in zebrafish notochord progenitors, while the cell adhesion module was upregulated in both progenitors and vacuolated cells, the latter being absent in medaka at comparable stages (Fig. 5D). Molecular processes implicated in timing can differ substantially in their cell type-specific deployment across species, and these differences serve as potential entry points for understanding species-specific developmental timing control. Overall, these findings suggest that developmental rate is not governed by a small set of genes driving acceleration or delay on demand and instead reinforces the idea that developmental rate modulation is constrained by existing cell type-specific functions.

## Discussion

Heterochrony can give rise to phenotypic diversity across species, but before its mechanistic basis and genetic drivers can be uncovered, we need a quantitative understanding of its prevalence and relationship to organism-level developmental tempo. We provide a time-resolved, cross-species data resource and a generalizable tool for systematic mapping of heterochrony between species using single-cell genomics. In comparing developmental progression at both the embryo- and cell-level, heterochrony can be defined as a deviation from embryo-wide progression, revealing widespread variation not predicted from other methods. Complementing morphological and imaging-based approaches (*22*, *28*, *51*), our single-cell transcriptomics approach increases sensitivity of developmental timing comparisons and provides a shared timescale to align additional morphological and molecular information derived from other methods.

Our approach revealed new insights into the relationship between cellular function and developmental timing in two ways: (1) in functional elaboration of a conserved cell identity, and (2) in defining species-specific phenotype by changing relative proportions of cells. In medaka, we find that ECM and calcium signaling components of a transcriptional program associated with later stages is deployed in the context of the early epidermal lineage, imbuing these cells with a species-specific contractile function during key developmental stages of epiboly (*32*, *52*, *53*). Thus, cross-species comparison of embryogenesis benefits from a wide temporal scope; emergence of novel cellular function via temporal redeployment of existing programs may contribute to the diversification of early developmental events occurring in a few, conserved cell identities (*54*). We find that the morphology of the notochord in medaka depends on a dual heterochrony that would not be predicted from the overall slower development of this species relative to zebrafish. Although notochord progenitors are globally delayed relative to zebrafish, their differentiated descendants diverge: sheath cells exhibit accelerated maturation while vacuolated cells show delayed maturation. Thus, developmental timing differences can shape tissues without invoking new cell identities, even when underlying cells share a lineal origin, revealing wider potential for timing variation than previously appreciated. Beyond describing cells as “faster” or “slower” in one species relative to another, or as a uniform stretching of development (*55*), our data indicate that relative timing differences can change in sign and magnitude as development proceeds, suggesting that timing control mechanisms must be capable of fine tuning across cells and over the course of development.

In systematically measuring timing variation across all cell types in the embryo, we showed that medaka’s slower developmental rate does not direct a uniform delay to all cells in the embryo. Our results challenge the assumption that species-level differences in developmental tempo arise from proportional temporal scaling of constituent cell types, with important implications for the relationship between *in vitro* cellular dynamics and those predicted at the level of the organism (*5*, *7*, *19*). Between species, we see a surprising lack of constraint in coordinated developmental progression at the cell-level, a trend that would be difficult or impossible to infer from isolated cell populations or morphological staging alone. We propose that organism-level developmental tempo may be better understood as an emergent property of underlying cell-level dynamics rather than as a uniformly imposed rule; that is, embryo development progresses faster when its cells progress faster. In this view, the limits of developmental tempo are not set by a global clock mechanism, but by the degree of cellular coordination, or synchrony, required to reproducibly build a viable embryo. For example, further delay of notochord progression relative to other cells may be limited by an essential signaling role that ensures embryo-wide progression. Our model also predicts that embryo-level developmental tempo could be slowed or stopped by the de-synchronization of one or a few cell types, so long as viability is maintained. Testing these predictions will require analysis of further species and genetically diverse populations to understand how cell type-specific timing differences influence phenotypic variation.

How do known developmental tempo control mechanisms operate across cell types? Among cells showing a relative delay, we identified processes associated with core gene expression and metabolism. Among cells showing relative acceleration, we identified processes linked to cell movement and morphogenesis, suggesting that physical cell interactions and tissue migration may contribute to developmental timing variation. Epigenetic and metabolic control processes (e.g., oxidative phosphorylation and histone methyltransferase activity) emerged as broadly relevant across cell types, but, like many of the pathways we identified, they were not exclusively associated with faster or slower development, suggesting that the effects of even broadly-expressed control mechanisms depend on cellular context. Across all trajectories examined, genes associated with acceleration and delay were enriched for conserved 1-to-1 orthologs, indicating that developmental timing variation arises from common and conserved molecular pathways rather than species-specific genes.

Rather than enforcing precise adherence to temporal patterns, developing embryos may tolerate or even benefit from flexibility in the timing of cell type emergence and maturation. This flexibility may contribute to developmental robustness in the face of environmental, chemical, or genetic perturbations that affect the timing or maturation of specific cell populations. On an evolutionary timescale, relaxing the requirements for temporal coordination across cells may provide additional degrees of freedom in the program of development, allowing cell types to independently shift timing without triggering failure of development embryo-wide. The broad thermal tolerance of medaka suggests an evolutionary history that required adaptation to fluctuating environments, while zebrafish develop rapidly and display a narrower thermal range, potentially indicating species-specific differences in the degree of cell-level timing variation. Teleost fish are the most diverse vertebrate lineage, with several groups, medaka included, showing recent phenotypic radiations. Developmental timing differences among species, strains and populations may provide substrate for adaptation and explain how embryogenesis tempers the mapping of genotype-to-phenotype.

## Acknowledgements

We thank the National BioResource Project medaka for providing medaka Hatching Enzyme. We thank Felix Loosli (KIT) for providing medaka HNI embryos, and Héctor Sánchez-Iranzo (KIT) for generously providing the Tg(*jag1a::mNeonGreen*) zebrafish line used in this study. We also thank members of the Centanin, Wittbrodt, and Dorrity labs for assisting in cell type annotation.

## Author Contributions

S.J.K, J.J.B, S.A.K, and M.W.D conceived and designed the study; S.A.K. optimized nuclei extraction and embryo hashing with input from L.M.S. and M.W.D.; S.A.K. collected samples with support from J.B., L.Z. and performed all sci-RNA-seq.; J.J.B. curated data and performed all computational analyses with insights from M.W.D.; J.J.B., S.A.K. performed live and *in situ* imaging experiments with support from S.J.K.; S.A.K. performed early medaka embryo HCR stains.; M.W.D. supervised the project; J.J.B. and M.W.D. interpreted the results and wrote the manuscript with input from all co-authors.

## Data, code, and materials availability

The raw sequencing data generated and the processed FASTQ files used in this study have been deposited in the European Nucleotide Archive (ENA) at EMBL-EBI (accession number pending). Raw image datasets and the corresponding metadata are deposited in the BioImage Archive at EMBL-EBI under (accession number pending). Processed single-nucleus datasets and a user-friendly data browser will be made available at our website: <pending url>. All original code, including notebooks for performing data processing, statistical analysis and generating plots, has been deposited at Github and will be made publicly available as of the date of publication: <pending url>.

## Supplementary Materials

Materials and Methods

Figs. S1 to S5

Tables S1 to S5

Movies S1 to S2

## Materials & Methods

### Animal rearing

Medaka (*Oryzias latipes*) strains used in this study were: Cab, Kaga, HNI, MIKK 72-2 and MIKK 79-2. Embryos for single cell transcriptomics were obtained from adults raised at the Centre for Organismal Studies (COS), Heidelberg (Cab, Kaga, MIKK 72-2 and MIKK 79-2) or the Karlsruhe Institute of Technology (KIT), Karlsruhe (HNI). Cab embryos for HCR and live confocal imaging were obtained from adults raised at EMBL, Heidelberg. All adult medaka fish were maintained at 26 °C under 14 h:10 h light:dark cycles. Zebrafish (*Danio rerio*) stocks used in this study were obtained from AB2B2 adults raised at EMBL, Heidelberg.

Zebrafish adult fish were maintained at 28.5 °C under 14 h:10 h light:dark cycles. Medaka and zebrafish embryos were raised at 26 °C in ERM and 28 °C in E3 medium, respectively, and sampled at indicated timepoints between 6 hpf and 216 hpf. Staging was performed according to Iwamatsu (*39*) (medaka) or Kimmel et al. (*40*) (zebrafish). All animal experiments were carried out according to the guidelines of the Committee for Animal Welfare and Institutional Animal Care and Use (IACUC) under EMBL’s Policy on the Protection and Welfare of Animals Used for Scientific purposes (IACUC code 22-003_HD_MD).

### Preparation of nuclei from barcoded individuals

Medaka embryos between 6.5 and 216 hpf were hatched using hatching enzyme (NBRP) two hours prior to dissociation. Nuclei isolations were performed as in previous zebrafish sci-RNA-seq (*25*) with modifications summarized here (see also Supplementary Protocol).

Embryos older than 24 hpf were transferred to individual wells of a 96-well plate for dissociation as described, but for embryos younger than 24 hpf, four embryos were transferred per individual well. Fixation was performed with the amine-crosslinker, 3,3’-dithiobis(sulfosuccinimidyl propionate) (DTSSP) dissolved in methanol. To affix “hash” oligos to nuclei during DTSSP fixation, ssDNA oligos were aminylated using a Label-IT nucleic acid modifying kit (Mirus Bio, cat. no. MIR-3925). Nuclei were then rehydrated and washed in 0.3 M SPBSTM 1X dPBS, 0.3M sucrose, 10% (v/v) Triton X-100, 3 mM MgCl_2_). Samples were frozen in 0.3 M SPBSTM using a FreezeCell™ Cell Freezing Container (Genesee Scientific, 27-802) and stored at - 70 °C. A step-by-step protocol detailing the preparation of aminylated ‘hash’ oligos and alterations to the nuclei isolation procedure can be found in the Supplementary Materials.

### Construction of sci-RNA-seq3 libraries

The nuclei were processed using the sci-RNA-seq3 protocol (*24*). Briefly, fixed nuclei were thawed and then spun down at 700 *g* for 3 min, resuspended in 400 μL 0.3 M SPBSTM. The nuclei were sonicated on low speed for 12 s in a Bioruptor Plus sonication device (Diagenode) and then spun at 700 *g* for 3 min. For the first barcoding step, 4 million nuclei were mixed with dTNPs (NEB, N0447L) to a final concentration of 1 mM and distributed evenly across the wells of two Lo-bind 96-well plates (Eppendorf, 30129504). Next, 2 μL of 10 mM 3-level-RT primers (IDT) containing the first index were added to each well of the plates. The plates were incubated at 55 °C for 5 minutes and SuperScript™ IV Reverse Transcriptase (Sigma Aldrich, 18090050) mix was added. The plates were then incubated in a thermocycler and subjected to 2-minute cycles of 4°C, 10°C, 20°C, 30°C, 40°C, 50°C before being held at 55°C for 15 minutes. Cold 0.3 M SPBSTM was added and all wells of both plates were then pooled. The pooled sample was spun at 700 *g* for 3 minutes and the pellet washed with 0.3 M SPBSTM. After spinning again, the pellet was resuspended in 0.3 M SPBSTM and 11 μL was distributed to each well of two fresh 96-well Lo-bind plates. The second index was then introduced by adding 2 μL of 10 mM 3-level-ligation primers (IDT) to each well of each plate. Ligation was performed by adding T4 DNA ligase in 1X T4 ligation buffer (NEB, M0202L) and incubating the plates at room temperature for 20 minutes.

Following this reaction, cold 0.3M SPBSTM was added to each reaction before all wells of the two plates were pooled. The sample was spun down and washed with 0.3M SPBSTM as described above three times and then counted using C-Chip counting chambers (Carl Roth, PK36.1). A total of 600,000 cells were then isolated, spun down as previously described, and resuspended in 500 μL NEBNext® Ultra™ II Non-Directional RNA Second Strand Synthesis mix (NEB, E6111L). 5 μL of the nuclei/second strand enzyme mix was then added to each well of a 96-well plate (Eppendorf, 30128656) and incubated at 16 °C for 2.5 hours followed by 4°C overnight.

The following day, DNA was released from the nuclei by adding 1 μL protease (Qiagen, 19157) to each well and incubating the plate at 37 °C for 30 minutes. The protease was then inactivated by incubating the plate at 75 °C for 25 minutes. Four wells of the 8-well PCR strip tube were then segregated to perform a Tn5 transposase dilution series test. An initial dilution of 0.05 mg/mL of Tn5 (purified EMBL stock) pre-loaded with annealed N7 Oligo (Eurofins) was added to TD buffer (20 mM Tris-HCl [pH 7.6], 10 mM MgCl_2_, 20% dimethylformamide) and then serially diluted 1:2 four times. Tagmentation was performed by adding 5 μL of each dilution of Tn5 mix to 5 μL of nuclei and incubating the samples for 5 minutes at 55°C. The reaction was stopped by adding transpose stop buffer (0.15 % SDS, BSA) and incubating at 55°C for a further 15 minutes. The reaction was quenched with 10% Tween-20. A 16-cycle PCR (70°C, 3 minutes; 98°C, 30 seconds; 16 cycles of 98°C for 30 seconds, 63°C for 10 seconds and 72°C for 60 seconds; 72°C, 5 minutes) was then performed on each dilution test using NEBNext high fidelity 2X PCR master mix (NEB, M0541L), indexed TruSeq P5 primers and indexed Nextera P7 primers (IDT). Post-PCR, samples were cleaned up with 1X AMPure XP Beads (Beckman Coulter, A63881). Briefly, beads were mixed and added to a concentration of 1X. After resting for a minute, tubes were added to a magnetic plate and the supernatant was removed. Following two washes with 80% ethanol, samples were air dried and then eluted in an elution buffer (EB; 10 mM Tris pH 8.5). Each sample was then analyzed on a Tapestation 4200 system (Agilent). The best concentration of Tn5 transposase was selected based on the proportion of the library between 250-700 base pairs. The remainder of the main plate was then processed with the chosen Tn5 concentration through tagmentation, stopping of tagmentation, quenching, PCR and tapestation analysis as above. This protocol was run twice for medaka nuclei from 26 - 216 hpf embryos and once for medaka nuclei from 6.5 – 18 hpf. For these younger embryos, only 96 indexed 3-level-RT primers were used and 220,000 cells were loaded into the PCR plate.

### Sequencing, read processing and cell filtering

Sequencing was performed on an Illumina NextSeq2000 (P2 kits, 100 cycle kit) and an Illumina Novaseq 6000 (S4 200 cycle kit) with the following sequencing chemistry: Index 1, 10 bp; Index 2, 10 bp; Read 1, 34 bp; Read 2, remaining cycles. Read alignment and gene count matrix generation was performed using the Brotman Baty Institute (BBI) pipeline for sci-RNA-seq3 (https://github.com/bbi-lab/bbi-sci). Sequences were aligned to NCBI *Oryzias latipes* annotation release 103 ASM223467v1. After the single cell gene count matrix was generated, cells with fewer than 150 (MA.strains), 400 (Olat.MA1) or 1000 (Olat.MAe) unique molecular identifiers (UMIs) were filtered out. Cells with a Scrublet score of ≥ 0.3 were also filtered out. For mitochondrial signatures, we aggregated all reads from the mitochondrial chromosome and removed all cells with ≥ 25% mitochondrial reads.

Background loadings for Olat.MA1 and Olat.MAe were calculated using “cells” identified during read processing as having between 3 and 20 UMIs. Each cell was assigned to a specific medaka embryo based on the total hash count (≥ 5) and the enrichment of a particular hash oligo (≥ 3).

### scRNA-seq analysis and major group clustering

After RNA and hash-quality filtering, data were processed using the Monocle3 (v.1.3.7) workflow defaults except where specified: *estimate_size_factors()*, *detect_genes()*, *preprocess_cds(num_dim = 100)*, *align_cds(residual_model_formula_str = “∼log.n.umi + bg_loading_MA1 + bg_loading_MAe”)*, *reduce_dimension(max_components = 3, umap.n_neighbors = 50L, pre- process_method = “Aligned”)* and finally, *cluster_cells(resolution = 5e-7)*. Clusters with fewer than 100 cells (17 in total) were removed from the dataset and the remaining 32 clusters were assigned to one of 14 major groups - roughly representing tissue groups (e.g. muscle, retina, liver) comprising all timepoints, and a single major group containing cells only from the earliest timepoints (6.5 hpf to 26 hpf), representing the blastula to late neurula stages - based on marker genes identified using the *top_markers()* function. Each cellular major group was reprocessed to identify and remove clusters with an overrepresentation of doublets and to create cluster annotations corresponding to cell types. Each of the major groups was first re-processed following a similar Monocle3 workflow as previously: *estimate_size_factors()*, *detect_genes()*, *preprocess_cds(num_dim = 50)* using only genes with expression variance ≥ 1.5 standard deviations from the mean of all genes in the major group, *align_cds(residual_model_formula_str = “∼log.n.umi + bg_loading_MA1 + bg_loading_MAe”)*, *reduce_dimension(max_components = 3, umap.n_neighbors = 30L, pre-process_method = “Aligned”)* and finally, *cluster_cells()* using a resolution dependent on the number of cells in the major group (>200000 = 1e-05, >150000 = 5e-05, >80000 = 5e-04, >10000 = 5e-03, otherwise 1e-02). To identify doublet-enriched clusters, the average Scrublet score, average hash oligo enrichment and total cell count were calculated for each sub-cluster and any cluster with a Scrublet mean > 0.1 and either a mean hash oligo enrichment ≤ 3.5 or cell count ≤ 80, or both, was removed. Following this removal, the major groups were reprocessed using the same workflow as above and the final sub-cluster set was generated using *cluster_cells()* with resolution depending on major group size (>200000 cells = 1e-05, >80000 = 5e-05, >10000 = 1e-04, otherwise 1e-03). This resulted in a final count of 302 sub-clusters. Each sub-cluster was assigned a sub-cell type label based on marker genes, expression data of orthologs in zebrafish (downloaded from ZFIN), the mean age of cells, and knowledge from the literature.

### Embryo-level trajectory inference

Counts per cell type (‘cell_type_broad’) were summed per embryo to generate an embryo *x* cell type matrix whose counts were normalized according to cell recovery per embryo. We performed a multinomial fit using vglm(family = ‘multinomial’) in R accounting for the temporal effects on cell composition with model formula ‘∼sm.ns(timepoint, df =3)’. With this model, we emitted predicted cell compositions for medaka embryos at 30 minute time resolution. We reduced dimensions of the 278 x 499 embryo matrix, containing both real embryo samples and dummy predicted embryos, using PCA and visualized the embryo trajectory in the first three dimensions. For the cross-species case, we performed the same procedure starting from the co-embedded medaka and zebrafish dataset, grouping cells from both species using the transferred ‘cell_type_broad’ label from zebrafish.

### Co-embedding of zebrafish and medaka strain cells

The zebrafish sci-Plex dataset was downloaded from https://cole-trapnell-lab.github.io/lmx1b/downloads/. The zebrafish atlas was subsampled to evenly represent the dataset’s sampled timepoints, and the medaka atlas was filtered to only include hashed cells from embryos 34 hpf or older. This resulted in a total of 867 652 medaka and zebrafish cells combined. The conserved 1:1 orthologous genes between zebrafish and medaka were downloaded from Ensembl BioMart (release 115) (*56*) and filtered for presence in both the zebrafish and medaka single-nucleus atlases, resulting in 15 011 shared gene features. The zebrafish and medaka datasets were then co-embedded using only the shared orthologous gene set. The co-embedding was performed using Monocle3, implementing nearest-neighbour-based batch correction across species: *estimate_size_factors()*, *detect_genes()*, *preprocess_cds(num_dim = 100)*, *align_cds(alignment_group = “species”, residual_model_formula_str = “∼log.n.umi”)*, *reduce_dimension(max_components = 3, umap.min_dist = 0.2, umap.n_neighbors = 30L, pre-process_method = “Aligned”)*. The medaka strain dataset was combined with 100 000 randomly chosen cells from the main (Cab only) medaka atlas. The strain and reference medaka cells were then co-embedded as before using Monocle3, implementing nearest-neighbour-based batch correction across strains: *estimate_size_factors()*, *detect_genes()*, *preprocess_cds(num_dim = 100)*, *align_cds(alignment_group = “genotype”, residual_model_formula_str = “∼log.n.umi”)*, *reduce_dimension(max_components = 3, umap.min_dist = 0.2, umap.n_neighbors = 30L, pre-process_method = “Aligned”)*.

### Cross-species assignment of cell type and developmental age

Nearest-neighbour label transfer was used to propagate developmental time and cell-type annotations both within and across species. Briefly, cells (either all medaka strains, or zebrafish and medaka Cab) were embedded in a shared UMAP space, and nearest-neighbour indices were constructed using cosine distance (*57*). For each query cell, the 20 nearest neighbours were identified in the reference population, and only neighbours within a fixed distance threshold (cosine distance ≤ 0.05) were retained to ensure local neighbourhood consistency. For continuous developmental age, the inferred value for each query cell was calculated as the mean sample timepoint of its neighbours, allowing us to assign each cell a “transcriptional age”, both within species and across species or strains.

Cell-level estimates were subsequently averaged across all cells belonging to the same embryo to obtain embryo-level transcriptional age values. Embryos with <200 cells were not assigned embryo-level ages. For categorical variables such as cell types, broad cell types and tissues, labels were assigned using a consensus-based approach. A label was transferred only when there was a consensus in more than 50% of the neighbours, or when the next-most frequent label was 1.5 times less enriched. This allowed the transfer of cell type labels from the medaka Cab cells to the other medaka strain cells, and from the zebrafish cells to the medaka Cab cells.

### Extracting cell type trajectories

Cell type-specific trajectories were analyzed separately for zebrafish and medaka using the species-specific atlases. For each species, only cells originating from hashed embryos were retained. Corresponding cell types were identified based on transferred and manually curated cell-type annotations shared across species. For each cell type, a corresponding subset of cells was extracted and reprocessed independently, using the previously described method for hierarchical cell clustering. Cells that lay strongly outside the cell trajectory or which lacked clear cross-species counterparts were removed, and processing was repeated on the filtered data to obtain a final trajectory for each cell type in both species. To confirm cross-species comparability, cell subsets were examined in the integrated cross-species embedding to verify spatial overlap or high proximity in the integrated space. Marker gene expression patterns were then examined for each trajectory and major sub-populations to ensure matched cell identity between species.

### Calculating CED values and across- and within-species

A within-species CED (“null”) distribution was generated for every hashed cell in the zebrafish sci-Plex dataset by subtracting every cell’s whole-embryo age from its developmental (“transcriptional”) age. The within-species CED distribution for hashed medaka cells was generated similarly, but using the transferred zebrafish cellular and whole-embryo age values. CED values for each cell in both species were then calculated per cell as CED = mean_nn_time − mean_nn_stage. CED values were calculated for *tbx16* mutants from published Zscape atlas (*25*) using the same approach, subtracting mutant embryo age (mean_nn_stage) from mutant cell age (mean_nn_time). To compute false discovery rate for medaka-specific CED values in the notochord, we used within-species variation of zebrafish notochord CED values as a null distribution. Using a smoothed density estimate from the null distribution, we used proportions of CED values in the test vs null distribution to estimate a false positive rate for both negative and positive values of CED, which was in turn used to compute false discovery rate.

### Correlating expression of medaka and zebrafish 1:1 orthologs in the notochord

Gene expression dynamics were compared across species within the notochord using embryo-level notochord time in zebrafish hours in both species. For each species, embryos with ≥5 notochord cells were retained and an embryo-level notochord stage was computed as the mean nearest-neighbour zebrafish time within the notochord. Notochord-expressed genes were filtered to those detected in ≥3% of notochord cells and restricted to conserved 1:1 orthologs. For each ortholog, expression was modelled as a function of notochord age using Monocle3 *fit_models()* with a natural spline (*splines::ns(mean_nn_zf_ct_stage, df = 3)*), and fitted values were obtained using the *model_predictions()* function. Predicted expression values were max-normalized per gene separately in each species and averaged in 1-hour bins. The similarity of the zebrafish and medaka temporal pattern of expression for each gene was quantified by Pearson correlation of the binned fitted expression profiles.

Peak expression time (the zebrafish hour in which each gene displayed its normalized maximum fitted expression) was computed for each gene in each species. Thresholds for the most conserved and diverged genes were defined using the 15th and 85th percentiles of temporal correlation.

### Hybridization chain reaction (HCR) preparation and mounting

Hairpins and buffers for HCR were purchased from Molecular Instruments (MI; Los Angeles, CA, USA). HCR probes were generated using a hybridization chain reaction probe generator tool (*58*) and ordered from IDT as Oligo Pools. One hour prior to fixation, zebrafish embryos were manually dechorionated with forceps and medaka embryos were hatched using medaka hatching enzyme (NBRP medaka). Embryos were staged (*39*, *40*) at indicated time points and then anaesthetized with MS222 (Sigma). Zebrafish and medaka embryos were then fixed overnight at 4°C with 4% PFA in 1X PBS or in 2X PTw (2X PBS, 0.1% Tween 20), respectively. Samples were then titrated into 100% methanol via 10 minute incubations with 1X PTw and then 25%, 50% and 75% methanol in 1X PTw. Fixed embryos were stored at - 20°C for up to two weeks. HCRs were performed according to Bruce et al. (*59*). Briefly, samples were first titrated back into 1X PTw via 10 minute incubations with 75%, 50% and 25% methanol in PTw. Four washes with 1 X PTw were performed and then the embryos were permeabilized with detergent solution (50 mM Tris-HCl [pH 7.5], 1% SDS, 0.5% Tween 20, 1 mM EDTA [pH 8.0], 150 mM NaCl) for 30 minutes. Samples were pre-hybridized with probe hybridisation buffer (MI) for 30 minutes at 37°C. 1.6 pmol of each probe was then added directly to the sample and incubated at 37°C overnight. The following day the probe solution was removed and embryos were washed four times with probe wash buffer (MI) at 37°C and then twice with 5X SSCT (5X saline sodium citrate, 0.1% Tween 20) at room temperature. Samples were then pre-amplified with amplification buffer (MI) for 30 minutes at room temperature. Hairpin solutions were prepared by adding 6 μM of h1 and h2 of each hairpin pair to amplification buffer, incubating at 95°C for 90 seconds and then cooling to room temperature in the dark over a 30 minute period. After pre-amplification, the amplification buffer in the samples was replaced with the hairpin containing solution and incubated in the dark at room temperature overnight. On the third day, the hairpin solution was removed and samples were washed five times with 5X SSCT and then incubated with 1X PBS containing 1 μg/mL DAPI for 30 minutes in the dark. Samples were then either stored in 70% glycerol in 1X PBS, or directly mounted in the same medium for imaging.

Samples from 12 hpf for zebrafish and 34 hpf for medaka were manually deyolked using sharp forceps inside a glass dish containing 70% glycerol in 1 X PBS; all other samples were imaged with yolk still intact. Samples were then transferred onto a glass microscopy slide, oriented inside a drop of 70% glycerol in 1 X PBS using an eyelash tool, and covered with a glass coverslip. Edges were sealed using nail polish, which was allowed to air-dry for at least 30 minutes before being transferred to the microscope.

### Live imaging sample preparation and mounting

Live Tg(*desmog::eGFP*) or mScarlet-PCNA medaka (*47*, *60*) and Tg(*jag1a::mNeonGreen*) zebrafish (*61*) embryos were dechorionated enzymatically or manually, respectively, and anesthetized in 1X Tricaine in ERM or E3 medium. Embryos were then transferred individually into low-melting-point agarose (0.3% for zebrafish, 0.8% for medaka) within wells of an eight-well glass-bottom dish and oriented using an eyelash tool. Agarose was then allowed to solidify and covered with 1X Tricaine in embryo medium prior to microscopy.

### Microscopy

Live imaging of medaka embryo contractions was performed on a Nikon Ti-2 CSU W1 SORA Spinning disk confocal microscope using a P-Apo Lambda S 10X/0.45NA objective, maintaining a constant sample temperature of 22°C. Notochord HCR-stained samples were imaged on a Leica STELLARIS 8 laser-scanning confocal microscope using a HC PL APO CS2 20X/0.75NA objective. Notochord live imaging was performed on a Zeiss LSM880 laser-scanning confocal microscope using a Plan-Apochromat 20X/0.8NA M27 objective, inside a custom incubator setup maintaining a constant sample temperature of 27°C.

### Image analysis

All imaging data was processed and annotated using Fiji (ImageJ2 version 2.16.0) (*62*). PIV analysis was performed using MATLAB R2025a and PIVlab v3.12.001. Maximum-intensity Z projection timepoints of mScarlet-PCNA medaka embryo contraction live imaging were pre-processed using CLAHE with a window size of 64 pixels, and a highpass threshold filter Kernel size of 15 pixels within PIVlab. PIV was performed as Multipass FFT window deformation with a first pass of an interrogation area of 64 pixels and a second pass of 40 pixels. A heatmap of vector standard deviation over the course of 350 timepoints acquired at 10 second intervals, spanning the first 12 contractions of the embryo, was generated and exported within PIVlab.

For quantification of notochord HCR staining signal across the anteroposterior axis, a segmented line was manually fitted on maximum-intensity Z projections, along the entire notochord length of each sample labelled by the three marker genes, and quantification of length and intensity of each individual channel was performed. The spatial positions of intensity values were then normalized on a per-sample basis, such that intensity values spanned an arbitrary distance of 0 to 1 along the anteroposterior axis. To allow for comparison across timepoints within one species, all intensity values for a given gene were then normalized to the minimum and maximum intensity values across all samples of one species, such that the intensities spanned arbitrary units of 0 to 100.

For processing of notochord live imaging, raw image stacks were first separated by channel, and only the fluorescent channel was retained for analysis. For zebrafish datasets, stacks were first Z-projected using average intensity across informative Z-slices at all timepoints.

Time-series drift was corrected using rigid body transformation in StackReg (version 2.0.0). For better visualisation, image contrast was enhanced using stack-wide histogram normalisation (2% saturation). For medaka datasets, Z-slice alignment was performed prior to projection using cross-Z registration (rigid body transformation in StackReg), and stacks were trimmed to retain only informative Z-slices. Average intensity Z-projections were then generated, followed by time-series registration using StackReg. Contrast was enhanced using stack histogram normalisation (0.35% saturation).

### Cellpose model training

To train a segmentation model, representative frames were randomly sampled from the Z-projected image stacks across developmental stages and biological samples. Cells in each selected frame were manually annotated by drawing regions of interest (ROIs), generating a training dataset of 149 annotated images. These annotated images were used to train a custom segmentation model using the Cellpose-SAM (version 4.0.8) (*63*) graphical user interface (GUI) with default training parameters, producing a model optimized for segmentation of vacuolated cells in both medaka and zebrafish.

### Analysis of notochord live imaging

Z-projected image stacks were segmented in Fiji using Cellpose-SAM (2D) with the previously described custom-trained model and default parameters. Segmentation masks were exported and analyzed in R. For each frame, labelled segmentation images were used to extract cell area and centroid coordinates using EBImage (*64*). For each species, vacuolated cells were identified by filtering segmented objects based on size and position along the notochord midline to remove segmentation artefacts and non-vacuolated cells. Cell area measurements were then aggregated per timepoint across embryos and aligned on a common developmental timescale. Medaka developmental time was converted to the equivalent zebrafish time using a previously established staging model (*36*) fitted by LOESS regression.

### Differential expression analysis across CED scores

To identify genes associated with CED, differential expression was assessed using regression-based modelling implemented with Monocle3 *fit_models()*. Differential expression analyses were performed both within individual developmental trajectories and across all trajectories combined. For trajectory-specific analyses, each trajectory dataset was processed independently and genes expressed in at least 15% of cells were retained for testing. Expression was then modelled using the formula *∼ced_cell + developmental_epoch* (*developmental_epoch* = somitogenesis or organogenesis). For global analyses across trajectories, trajectory-specific datasets were combined and genes expressed in at least 5% of all cells were retained. Expression was modelled using *∼ced_cell + cell_type_broad + developmental_epoch*. Trajectory marker genes were additionally identified separately using *top_markers()* and strongly trajectory-associated genes (specificity >= 0.5) were removed.

### Gene ontology analysis

Differentially expressed genes associated with CED were classified based on the sign of their normalized regression coefficient. Genes with positive coefficients were defined as positive heterochronic regulators (“accelerating” genes), those with negative coefficients as negative heterochronic regulators (“delaying” genes), and genes appearing in both sets across trajectories were classified as bidirectional regulators. Genes were filtered to retain significant associations with CED (q < 0.05), and Gene Ontology (GO) enrichment analysis was subsequently performed separately for fast, slow, and bidirectional gene sets using medaka-specific GO annotations obtained from Ensembl BioMart. Enrichment testing was conducted using the clusterProfiler (*65*) *enricher()* function with Benjamini-Hochberg multiple testing correction.

### Gene module assembly

Gene modules were assembled from two sources: (i) GO-derived modules and (ii) DEG-derived modules. GO modules were generated by choosing a biologically relevant GO Biological Process term and expanding each term by obtaining all children of that term. All genes annotated to the resulting term sets were retrieved from the medaka Ensembl BioMart GO dataset and intersected with genes present in the atlas to create the final gene modules. DEG-derived modules were loaded as gene lists obtained from enrichment analyses of CED-associated genes.

### Obtaining per-trajectory module enrichment scores

Each of the 15 developmental trajectories was scored for enrichment of every module using Seurat *AddModuleScore(nbin = 20, ctrl = 50)*. For each trajectory, individual cells received a module score reflecting the aggregate expression of genes within that module. Module scores were z-scored across cells in all trajectories. Cells were considered “module-enriched” if their scaled module score exceeded ±2 standard deviations. For trajectory-level summaries, module-trajectory pairs were retained only if 5% of the cells in the trajectory exceeded the module score threshold. For each retained module-trajectory pair, we calculated the mean CED and mean module score among enriched cells.

**Fig. S1.**
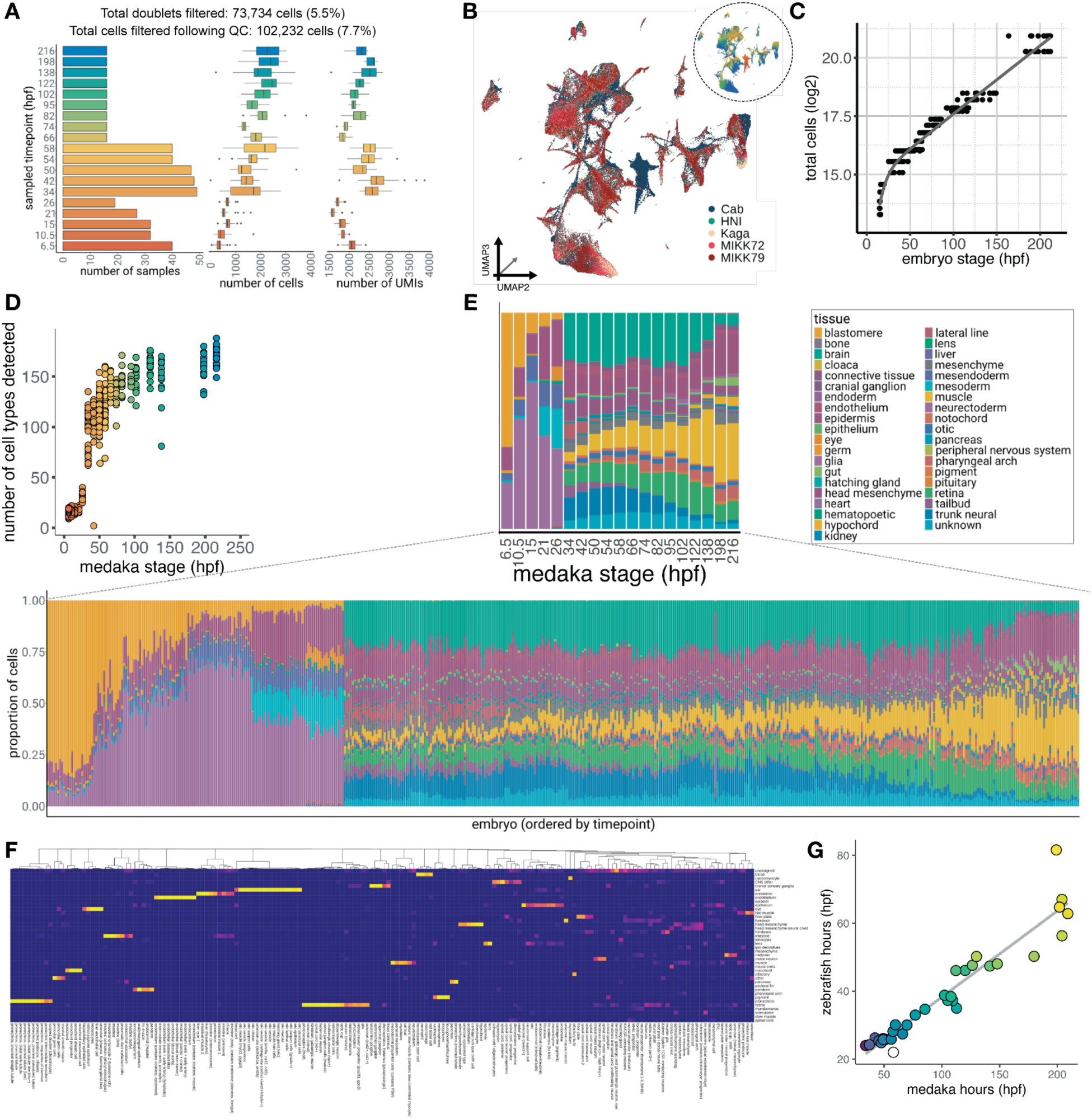
Quality control of medaka time-series atlas. (A) Sampling strategy and dataset composition across medaka development. Left, number of embryos profiled at each developmental timepoint (hours post fertilisation, hpf). Middle, the distribution of the number of nuclei recovered in all embryos per timepoint. Right, the distribution of detected unique molecular identifiers (UMIs) in all embryos per timepoint. (B) UMAP plot showing medaka nuclei obtained from all five sampled strains (Cab, Kaga, HNI, MIKK72 and MIKK79) co-embedded in three dimensions, showing developmental processes are well-captured across medaka genotypes (colors); inset shows per-cell developmental age. (C) Number of cells per-embryo estimated by DNA content over the course of medaka development. (D) Number of distinct cell types detected over the sampled timepoints, showing increasing cellular complexity over embryogenesis, and a saturation of complexity towards later stages. (E) Relative proportions of the major tissues across sampled medaka developmental timepoints, illustrating dynamic changes in tissue composition during embryogenesis. Top, all cells per timepoint summarized; bottom, tissue composition per embryo (n = 518), ordered by increasing sampled timepoint. (F) Confusion matrix showing the proportion of medaka cells within each broad cell type annotation that were assigned to each zebrafish tissue, obtained via nearest-neighbour label transfer within the co-embedded medaka-zebrafish single-nucleus dataset. (G) Linear relationship between medaka and zebrafish developmental time inferred from whole-embryo cell composition-based pseudostaging in PCA space.

**Fig. S2.**
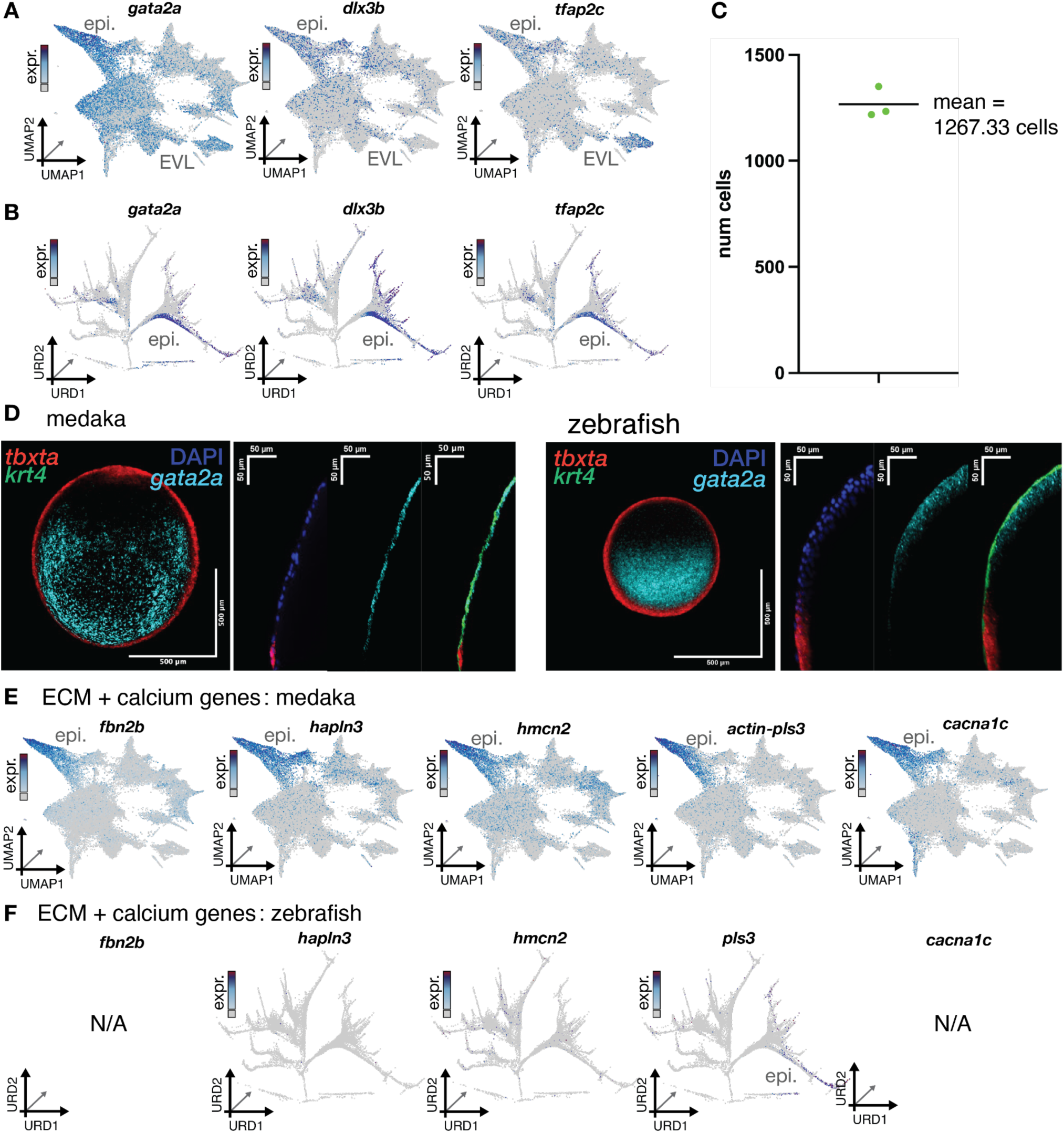
Medaka-specific contractile function in an epidermal cell identity shared with zebrafish. (A) UMAP plots of early medaka cells colored by log10 expression of non-neural ectoderm and early epidermis markers *gata2*, *dlx3b*, and *tfap2c*. (B) Marker gene plots for the same set of genes in A, shown in zebrafish (*32*). (C) Boxplot showing quantification of total *vcana*+ cells per medaka stained as in Figure 2D (n = 3). (D) HCR stains as in 2D showing expression of an additional epidermal marker, *gata2a*, in medaka (left) and zebrafish (right) confirming this cell population is present in both species. (E) UMAP marker gene plots for medaka showing expression of a set of ECM and calcium signaling genes (*fbn2b, hpln3, hmcn2, actin-pls3, cacna1c*) associated with contractile function and enriched in the medaka-specific stellate cell population. (F) UMAP marker gene plots for genes in E showing no expression in the epidermal population of zebrafish (see B above for reference); genes with no detected expression are labelled NA.

**Fig. S3.**
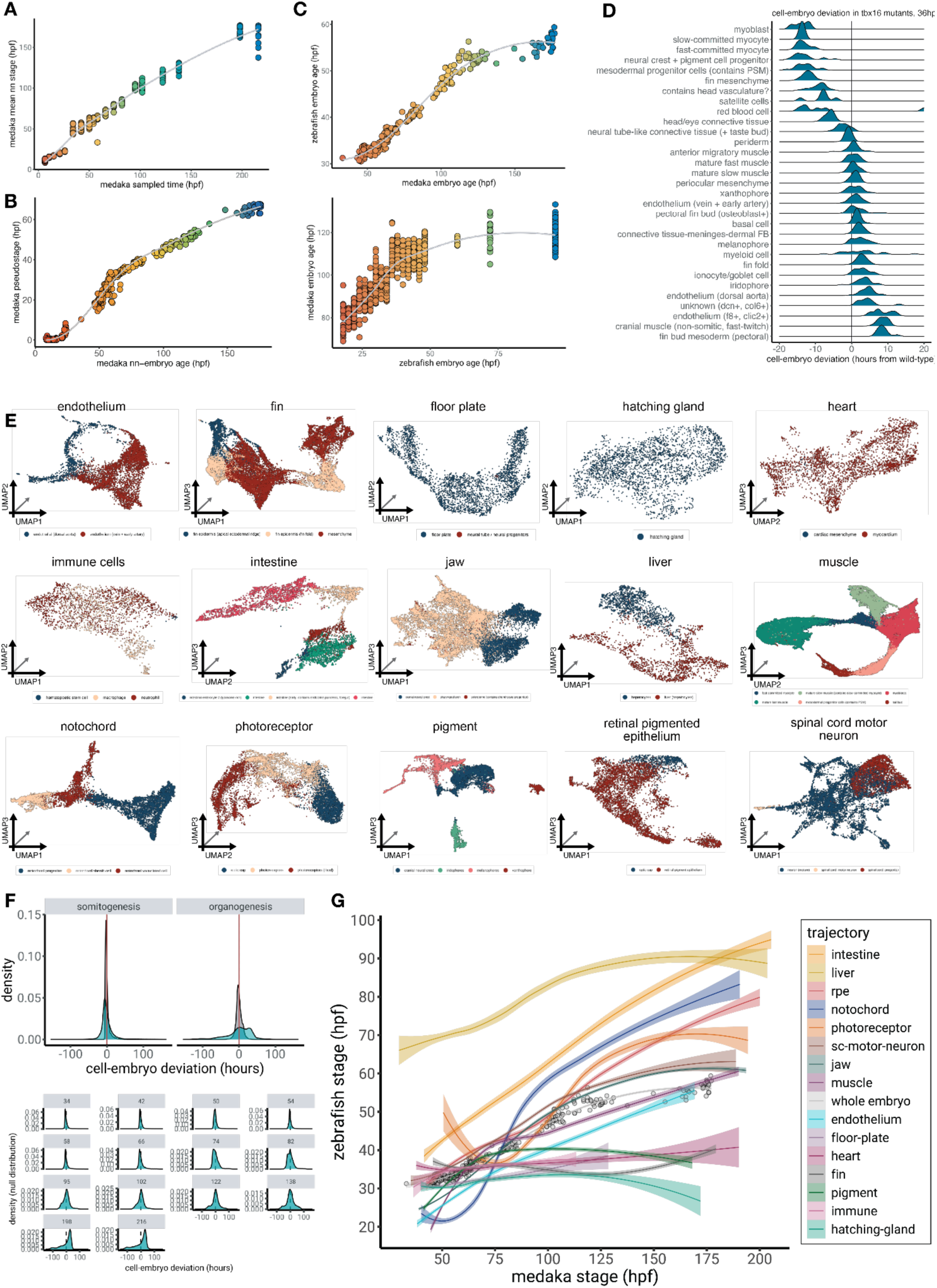
Quantitative developmental staging of embryos and cells. (A) Relationship between medaka whole-embryo sampled timepoint and medaka whole-embryo developmental age obtained via nearest-neighbour label transfer in the medaka atlas, showing a strong positive correlation. (B) Relationship between inferred medaka whole-embryo developmental age obtained via nearest-neighbour label transfer and inferred medaka whole-embryo developmental age obtained via cell-type composition (Figure 1C). (C) Cross-species whole-embryo developmental time relationships obtained via nearest-neighbour label transfer within the co-embedded medaka-zebrafish atlas. Top, medaka developmental times represented in zebrafish hours. Bottom, zebrafish developmental times represented in medaka hours. (D) Distributions of CED values in broad cell types of *tbx16* mutant zebrafish (*25*); mesodermal cells displayed notably negative CED values, indicating delay in development while progression of other cells is intact. (E) UMAP plots, embedded in three dimensions, of each of the 15 medaka cell type trajectories used to examine cell-type-specific dynamics. Colors show broad cell type. (F) The distribution of within-zebrafish CED values in the zebrafish dataset (grey) representing the null distribution, overlaid with the distribution of the medaka CED values (blue). Top, distributions are shown separately for the developmental epochs of somitogenesis and organogenesis. Bottom, timepoint-specific distributions of medaka CED values are shown across increasing developmental timepoints, illustrating the progressive changes in timing deviations during embryogenesis. (G) Cross-species developmental trajectories for whole embryo and selected tissues as in Figure 2D, shown for all 15 groups. Lines represent smoothed fits for whole-embryo alignment (grey) and individual cell types (colors), built on data where each point represents the average age of that cell type in each embryo (n = 367). Shaded regions indicate the 95% confidence interval.

**Fig. S4.**
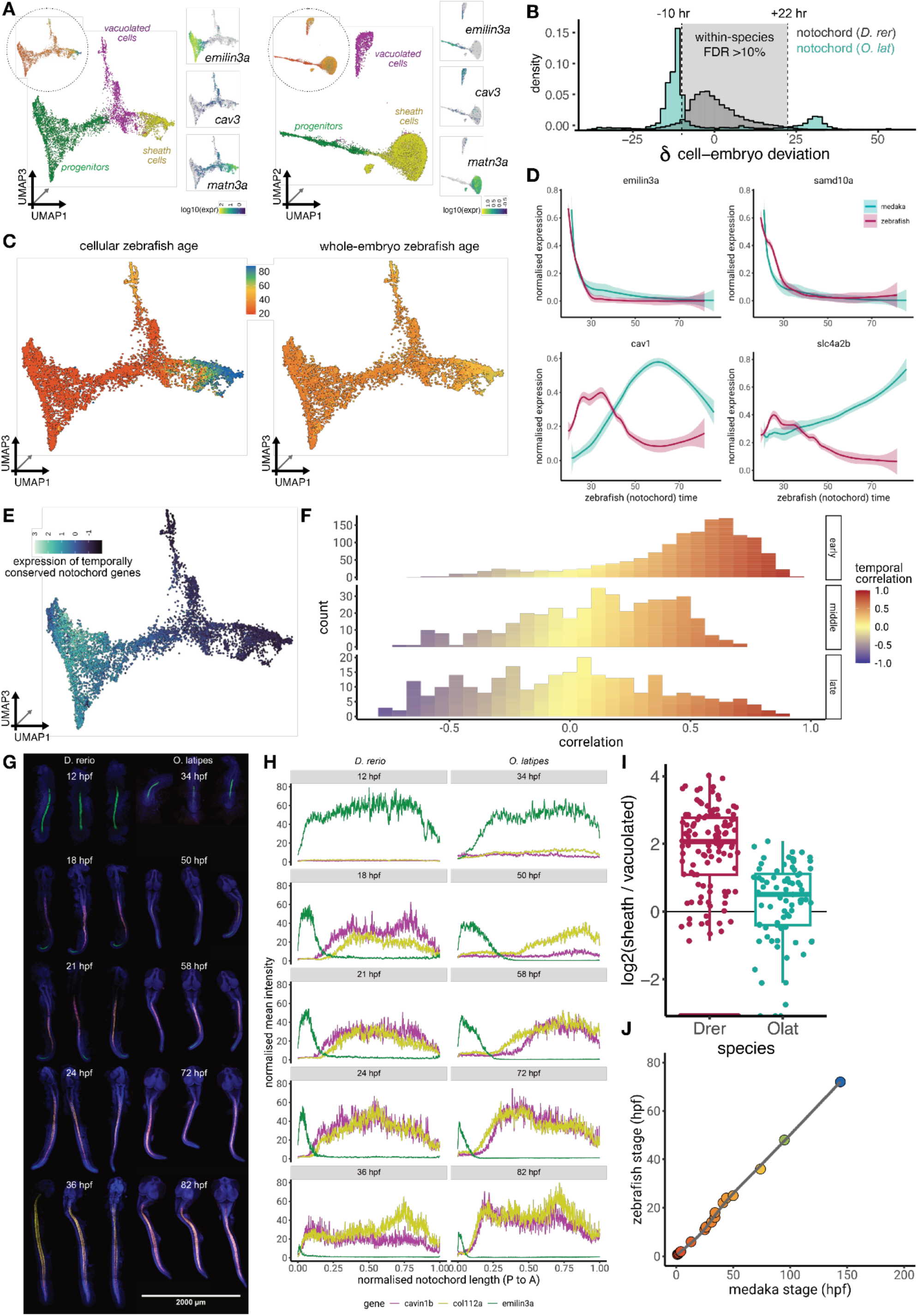
Medaka-specific heterochrony in two notochord cell sub-types. (A) UMAP plots of the medaka (left) and zebrafish (right) notochord cells embedded in three dimensions, colored by broad cell type, matching those represented in Figure 4A. Inset colored by per-cell developmental age. Marker plots for cell type-specific genes with 1:1 orthology are shown to the right of each species’ UMAP, colored by log10 expression. (B) Medaka notochord CED distribution (teal) as shown in Figure 2B, plotted alongside null distribution of within-species differences in zebrafish (grey), allowing computation of < 10% FDR thresholds for both delayed and accelerated cell populations. (C) UMAP plots of the medaka notochord cells embedded in three dimensions, colored by inferred zebrafish cell age (left) or inferred zebrafish whole-embryo age (right). (D) Representative examples of genes showing the most conserved (*emilin3a*, *samd10a*) and divergent (*cav1*, *slc4a2b*) modelled expression dynamics over time in the notochord trajectory of both species. (E) The aggregated expression of the 1:1 orthologous genes with the most conserved temporal dynamics across species. The progenitor population displays the highest expression levels of temporally conserved genes. (F) Distributions of temporal correlation coefficients of notochord-specific 1:1 orthologs across species, faceted by early (< 37 hpf), middle (between 37 and 71 hpf), and late (> 71 hpf) peak expression times in medaka. (G) Time-series of hybridisation chain reaction (HCR) stainings in whole embryo mounts, illustrating the progression of notochord cell differentiation in zebrafish (left) and medaka (right). Embryos are shown at morphologically stage-matched timepoints. Progenitor (green, *emilin3a*), differentiated vacuolated cell (magenta, *cavin1b*), and differentiated sheath cell (yellow, *col11a2*) populations are visualized, with nuclei stained with DAPI (blue), revealing species-specific timing differences in the onset and progression of notochord differentiation. n = 3 embryos for each species at each timepoint; scale bar = 2000 µm. (H) Quantification of cell type-specific HCR in G along the anterior-posterior axis (posterior = 0, anterior = 1) of the notochord, revealing medaka-specific pattern of sheath expression preceding vacuolated cell expression starting at the posterior. (I) Boxplots showing, for embryos >= 48hpf zebrafish time (fully vacuolated), the log2 ratio of sheath cell abundance to vacuolated cell abundance in each species. (J) Relationship between medaka developmental age (hpf) and corresponding zebrafish-equivalent age (hpf), obtained from morphological comparisons reported by Tena et al. (*36*).

**Fig. S5.**
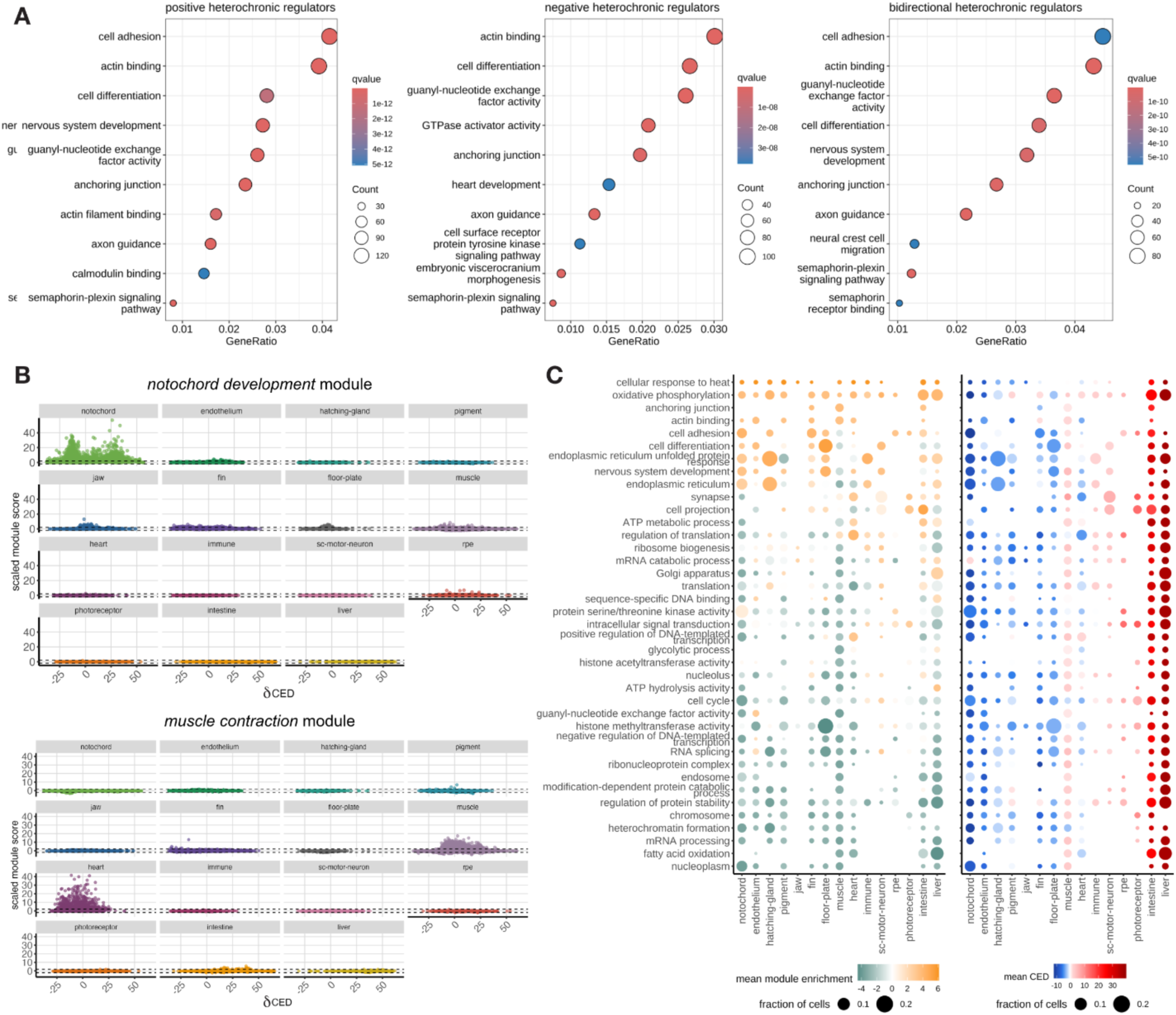
Molecular characterization of cell type-specific timing differences. (A) Gene Ontology (GO) enrichment analysis of all unique genes from 15 individual developmental trajectories analyzed that are significantly associated with acceleration (left; positive CED), delay (middle; negative CED), or both acceleration and delay of developmental timing in different cell types (right). Dots represent enriched GO terms, positioned by gene ratio, sized by the number of genes contributing to the term, and colored by adjusted q-value. (B) Distributions of scaled modules scores for test modules (top, notochord development; bottom, muscle contraction) plotted against cell-embryo deviation (CED) for each developmental trajectory. The dotted lines represent two standard deviations from the mean. Each control module displays enrichment in the expected cell types, confirming that the module construction approach accurately represents biological programs. (C) Summary of each module’s activity across developmental trajectories. Only trajectory-module combinations in which ≥2% of cells exceeded the two-standard-deviation enrichment threshold were included. Dot size indicates the fraction of cells within each trajectory showing significant module enrichment, while color represents mean module enrichment score (left) or mean CED (right).

**Table S1.**

Hierarchical cell type annotations (sub cell type, broad cell type, tissue, germ layer and major group) of each cluster in the medaka atlas.

**Table S2.**

All zebrafish and medaka orthologous genes, downloaded from Biomart (Ensembl release 115, Japanese medaka HdrR genome ASM223467v1) on 5 January 2026.

**Table S3.**

All 1-to-1 orthologs between zebrafish and medaka that were expressed in >= 3% of notochord cells within each species, their correlation values for modeled expression dynamics over zebrafish time, and the zebrafish time at which their expression reaches the peak value.

**Table S4.**

Globally (across all 15 trajectories) CED-associated genes, their normalized effect on CED and their test values following differential gene expression analysis in Monocle3.

**Table S5.**

All unique, trajectory-specific CED-associated genes, their normalized effect on CED and their test values following differential gene expression analysis in Monocle3.

**Movie S1.**

Particle image velocimetry (PIV) of a medaka embryo during gastrulation stages to quantify rhythmic contractile activity in the stellate cell layer.

**Movie S2.**

Confocal live imaging of a transgenic zebrafish and medaka through vacuolation of the notochord. Left, transgenic *jag1a::mNeonGreen* zebrafish embryo from 12 hpf to 30 hpf, played at a speed of 43 clocktime minutes per second. Right, transgenic *desmog::eGFP* medaka embryo from 34 hpf to 74 hpf, played at a speed of 95 clocktime minutes per second. The medaka movie was sped up by a factor of 2.22 to achieve the same movie timespan as the zebrafish, allowing comparison of the same vacuolation process in the same relative time.

